# SCOPE: Localizing fate-decision states and their regulatory drivers in single-cell differentiation

**DOI:** 10.64898/2026.04.07.717037

**Authors:** Yimin Zhao, Connor Finkbeiner, Manu Setty, Kevin Z. Lin

## Abstract

Identifying the precise transcriptomic states at which cells commit to a lineage (branchpoints) and the temporal lag in which chromatin accessibility foreshadows gene expression (epigenetic priming) remain fundamental challenges in developmental biology. While current methods for single-cell sequencing data effectively capture developmental flow, they often lack a principled mechanism for delineating the discrete boundaries, a crucial aspect required to map the molecular logic of lineage commitment. We present SCOPE (Semi-supervised Conformal Prediction), a framework that transforms high-dimensional single-cell measurements into rigorous, discrete prediction sets of all plausible future fates. By formalizing fate uncertainty via conformal inference, SCOPE localizes the precise biological windows during which multipotent progenitors specify their fate. In multi-omic data, SCOPE uncovers epigenetic priming and identifies its driving transcription factors by detecting regimes where chromatin-derived prediction sets resolve toward terminal fates significantly before their transcriptomic counterparts. We apply SCOPE across simulations, lineage-traced mouse hematopoiesis, multiple human hematopoietic datasets, and human retinogenesis to demonstrate its broad applicability and ability to recapitulate known fate specification drivers. Ultimately, SCOPE provides a statistically grounded foundation for localizing fate decisions across biological replicates and modalities, offering a robust tool for identifying the onset of lineage specification in complex developmental systems.

## 1 Introduction

As cells differentiate in pluripotent systems, they traverse intermediary states and gradually specify their fates to distinct terminal cell types. This transition is defined by two recurrent questions across developmental systems: First, where along a trajectory do cells reach the “decision state,” the branchpoint at which a cell is in a discrete transcriptomic checkpoint that precedes fate specification? Second, within this branchpoint, is there sufficient evidence of regulatory priming where chromatin becomes permissive for a lineage program before that program is fully expressed at the mRNA level [1, 2], and if so, where?

The precise state of cellular decision-making and the timing of TF regulatory drivers have remained difficult to localize, specifically, pinpointing the exact cell subsets where fate decisions occur and resolving when fate-specifying TF regulatory programs are activated. Resolving these transition states is essential for mapping the molecular logic that governs how multipotent progenitors exit their undifferentiated state and initiate lineage-specific programs. Current methods have made significant progress in modeling the differentiation continuum: pseudotime-based methods can order cells along a latent progression of transcriptomic change [3–8], while RNA velocity methods propose another perspective, utilizing the ratio of unspliced to spliced mRNA to infer the directional vector of individual cells [9, 10]. While these frameworks have revolutionized our mapping of developmental landscapes, translating continuous fate potential into discrete biological boundaries remains a challenge; yet, this resolution is essential for identifying the precise windows of plasticity where targeted interventions can be successfully deployed. For instance, in acute promyelocytic leukemia, the ability to pinpoint and block specific hematopoietic branchpoints has been shown to restore normal differentiation [11], highlighting how discrete boundaries serve as a necessary roadmap for therapeutic intervention.

To address the challenge of pinpointing these transitions, we develop SCOPE (Semi-supervised COnformal PrEdiction), a classification framework that leverages conformal inference [12–14] to compute the set of likely terminal fates a cell may become in the future. Unlike standard probabilistic models, these prediction sets provide a coverage guarantee: at a user-chosen error level *α* such as 0.05, the true terminal fate is guaranteed to be contained within the set with probability 1 − *α*. In developmental terms, this creates a rigorous, set-based language for “poise”: for hypothetical terminal fates *A* and *B*, a progenitor cell with a singleton prediction set (e.g., {*A*}, unipotent) indicates that a cell is already fate-specified, whereas a multi-fate prediction set (e.g., {*A, B*}, multipotent) indicates a cell that is still fate-neutral. This operational measure of fate uncertainty is interpretable, rigorous, and uniquely comparable across biological modalities, allowing us to localize the exact regimes of commitment. We then use these rigorously defined prediction sets to precisely localize where lineages branch and when the underlying regulatory programs are first activated.

We first demonstrate that SCOPE localizes developmental branchpoints when analyzing single-cell RNA-sequencing (scRNA-seq) data. Reinterpreting fate specification as a gradual loss of multipotency [15, 16], we define a branchpoint as a region of the cellular manifold where fate-neutral sets (e.g., {*A, B*}) are highly concentrated in transcriptomic space. By combining density-based clustering with conformal prediction, SCOPE targets the largest fate-neutral checkpoint immediately upstream of the commitment transition.

Localizing branchpoints does not, by itself, offer mechanistic insight into fate specification. Hence, when analyzing paired multi-omic single-cell RNA and ATAC data, we demonstrate that SCOPE’s prediction sets also quantify epigenetic priming [1, 10, 17– 19]. We hypothesize that if chromatin accessibility foreshadows gene expression, a cell’s ATAC-derived prediction set should collapse towards singletons (e.g., {*A*}) earlier in pseudotime, while its RNA-derived prediction set remains fate-neutral (e.g., {*A, B*}). This operationally defines a “priming window.” By pairing this with featureimportance analyses, SCOPE can nominate mechanistic candidates whose enhancers become predictive of fate before their target genes.

Applications across simulations, lineage-traced mouse hematopoiesis [20], and diverse human multi-omic datasets demonstrate that SCOPE reliably recovers biologically meaningful checkpoints and their molecular drivers. For example, in hematopoiesis, SCOPE allows us to recapitulate known specific TF regulatory drivers such as Gata2 and Klf4 that define the transcriptomic profile of the “decision-ready” state. Crucially, the framework yields consistent results across independent bonemarrow replicates [8, 21] and recapitulates known priming drivers in the mouse retina [22]. By establishing a statistically grounded tool for identifying fate-specification branchpoints, SCOPE enables researchers to connect molecular regulatory events to cellular decision-making across high-dimensional developmental landscapes, all within one unified statistical framework.

## 2 Results

### 2.1 Method overview

SCOPE localizes fate-decision states and their corresponding regulatory drivers from single-cell data by formalizing fate uncertainty through a semi-supervised conformal prediction framework (Fig. 1). Conformal inference provides the ideal framework for translating the inherent uncertainty of cellular decision-making into a rigorous, set-based language. Starting with labeled terminal cells and a user-chosen error level *α*, SCOPE models the fate specification process via a classification framework that computes each cell’s set of plausible terminal fates, thereby characterizing the uncertainty landscape across the entire transcriptomic space. Suppose {*X*_*i*_, *Y*_*i*_} represents a cell *i*’s gene expression profile and its (possibly unobserved) terminal fate, respectively. SCOPE uses conformal inference to construct a prediction set 𝒞 (*X*_*i*_) such that ℙ (*Y*_*i*_ ∈ 𝒞(*X*_*i*_)) ≥ 1 − *α* for a target error level *α* ∈ [0, 1], where typically *α* = 0.05. A defining advantage of conformal inference is that it is distribution-free; it requires no parametric assumptions regarding the underlying data distribution. Another key feature of this approach is that the primary tuning parameter, *α*, provides a concrete interpretation by explicitly controlling the target coverage probability and the size of the resulting prediction sets. These characteristics provide a robust, statistically grounded measure of “poise” that remains valid across diverse biological systems and sequencing modalities.

**Fig. 1:**
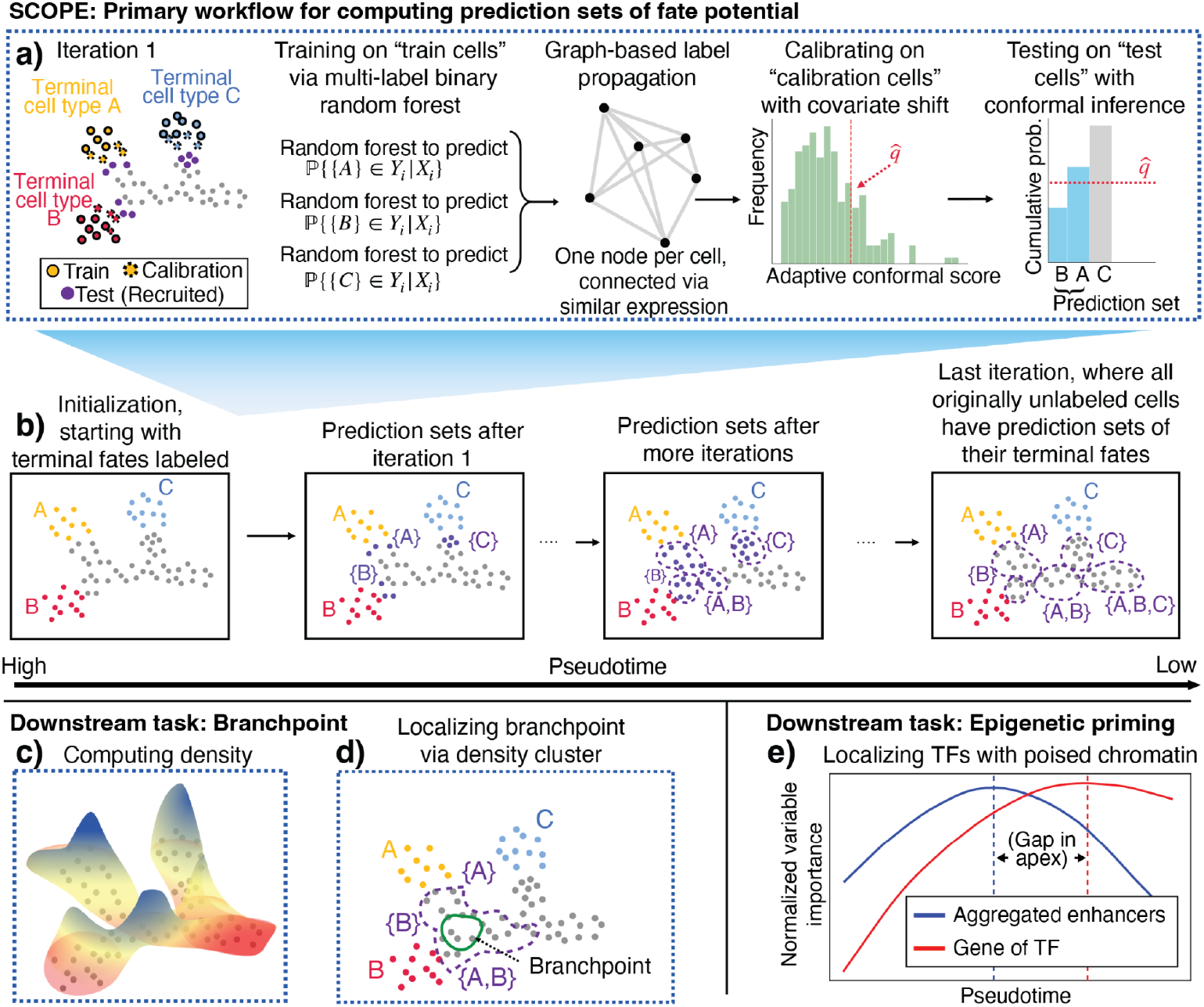
Schematic of the SCOPE model. **a**,**b)** SCOPE iteratively recruits cells along a pseudotime trajectory to generate calibrated prediction sets of terminal fates. This procedure involves training multi-label ensembles, smoothing predictions via graph-based label propagation, and applying weighted conformal inference to provide finite-sample coverage guarantees. **c**,**d)** Downstream application of branchpoint localization: SCOPE’s prediction sets are integrated with cell-state density landscapes to identify branchpoints where fate-neutral cells accumulate. **e)** Downstream application of epigenetic priming: SCOPE uncovers regulatory lags by contrasting importance scores between RNA and ATAC modalities for cells along the lineage, identifying genomic regions where chromatin accessibility becomes fate-predictive before gene expression.

At a high level, SCOPE starts with labeled terminal-state cells and iteratively trains classifiers while recruiting unlabeled cells along pseudotime. In each iteration, the labeled cells are split into training and calibration sets. We then train a binary-relevance ensemble (independent random-forest classifiers, one per fate) [23] on training cells. Then, for each cell in the calibration set, we predict its fate probabilities and compute the cumulative probability required to cover the cell’s labeled fate(s), ordered from most to least probable; this value defines the conformal score. The empirical *α*-quantile of these scores defines a threshold, 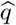, that determines the inclusion criteria for the prediction sets of the unlabeled cells. These newly formed prediction sets then serve as labels for the subsequent recruitment iteration. By repeating this process, SCOPE eventually generates calibrated prediction sets for all cells across the cellular manifold (Fig. 1a,b). We discuss SCOPE’s connection with the conformal inference in more detail in Supplementary Note S1.

To deploy conformal inference robustly across the developmental continuum, SCOPE incorporates two key technical advancements: label propagation and entropy balancing (see “Methods and analysis”). First, to improve stability in early iterations where labeled data are sparse, we use label propagation to smooth predictions across a *k*-nearest-neighbor graph [24]. This ensures that a cell’s predicted fate potential is transcriptomically coherent with its neighbors on the manifold. Second, because SCOPE recruits unlabeled cells along a trajectory, it must account for distribution shift, a statistical phenomenon where the transcriptomic profiles change significantly between the “mature” calibration cells and the “progenitor-like” test cells. To maintain accurate coverage despite this shift, SCOPE uses entropy balancing to reweight calibration cells so their conformal score distributions match those of the current test set [25]. This ensures that the statistical guarantees of the conformal framework remain valid as we move from terminal cell types back toward early progenitors.

To localize branchpoints after computing the prediction sets, SCOPE adopts a dynamic interpretation: high-density regions represent metastable regimes where cells accumulate, while low-density “valleys” signify rapid transitions in gene programs [9, 26]. Computationally, a prediction set of {*A, B*} identifies a cell as fate-neutral, but such cells may be distributed across disparate states or arise from transient noise. By defining a branchpoint as the intersection of high uncertainty and high cell-state density, we ensure that the identified population represents a coherent, metastable branchpoint immediately upstream of fate specification (Fig. 1c,d). This approach leverages density clustering to localize where fate-neutral cells physically converge on the manifold, providing a transcriptomically grounded definition of the “last state” before sets resolve into singletons, e.g., {*A*} or {*B*} .

Once these branchpoints are localized, SCOPE interrogates their molecular drivers by leveraging the feature importance scores from its underlying iterative classifiers. While conformal inference provides rigorous boundaries for defining these populations, the underlying random forests reveal the information value of specific features. Crucially, the SCOPE framework is agnostic to the molecular modality; the same predictive logic used for RNA can be applied to chromatin accessibility (single-cell ATAC-seq) or other high-dimensional biological measurements. This flexibility allows us to apply SCOPE across paired multi-omic data to directly contrast the timing of fate resolution and importance scores between the epigenome and the transcriptome. By identifying regimes where enhancer-based importance scores reach their apex significantly before their target genes (Fig. 1e), SCOPE captures regulatory priming events where chromatin accessibility becomes fate-predictive before the program is expressed at the mRNA level.

SCOPE’s design intentionally unifies developmental topology and regulatory timing by improving conceptual and statistical aspects of current methodologies. While trajectory-inference frameworks suggest global dynamics via fate probabilities, they lack a discrete transcriptomic definition of fate-specifying branchpoints; conversely, model-based methods often impose rigid topological priors that may obscure biological complexity (e.g., [4, 7, 9]). Similarly, current multi-omic approaches for epigenetic priming are often statistically disjoint from branchpoint localization and are often limited by expression-level thresholds sensitive to signal sparsity (e.g., [10, 18, 27]). See Supplementary Note S2 for an in-depth discussion on how SCOPE conceptually differs from other existing methods [28, 29]. To address these limitations, we deliberately use random forests as our fate classifiers rather than deep-learning architectures, as they provide performance stability for SCOPE’s iterative self-training regime in small-sample, imbalanced contexts (see Supplementary Note S3). By formalizing fate specification through calibrated prediction sets, SCOPE provides a mathematically principled language that consistently localizes both the cellular “states” of decision and the informational lag of the underlying transcription factor drivers across disparate biological modalities.

### 2.2 Simulated and lineage-traced scRNA-seq datasets both demonstrate coverage and power of SCOPE’s prediction sets

Before we assessed SCOPE’s ability to localize branchpoints or detect epigenetic priming, we first ensured that SCOPE estimated well-calibrated prediction sets. This aspect was critical, as these prediction sets underlie both biological tasks. We first interrogated this using simulated data with 240 genes and 800 cells, where we designed our own simulation suite to differentiation topology [30]. Existing single-cell simulators such as dyngen [31], scDesign3 [32], and scMultiSim [33] were not able to easily simulate data with complex trajectory topologies beyond a simple one-layer bifurcation (see Supplementary Note S8 for a comparison with dyngen). Our simulation sampled each cell’s gene expression from a two-dimensional latent space, which consisted of a branching structure among three terminal fates (Fig. 2a). SCOPE achieved above 95% coverage for each of the three terminal fates across multiple trials, and we further investigated SCOPE’s power through visualization. In Fig. 2b, cells highlighted in purple (cells whose prediction sets include the correct terminal fate) were concentrated along the branch leading toward terminal fate 1 (red). These cells were geometrically close to the labeled terminal cells in the PCA embedding and were clearly separated from the other two terminal fates. Fig. 2c,d revealed a second, downstream bifurcation. In Fig. 2c, the highlighted cells predominantly occupied the manifold leading toward terminal fate 2 (blue), whereas in Fig. 2d the highlighted cells instead aligned with the manifold leading toward terminal fate 3 (green). These two groups populated distinct branches of the manifold, demonstrating that after the initial split, the trajectory bifurcates again, separating cells destined for terminal fates 2 and 3. Taken together, we showed that the simulated system exhibits a hierarchical branching structure with two bifurcations in fate. We also compare SCOPE against FateID [34] in Fig. S2, which had limited performance in our simulated data due to its recruitment procedure.

**Fig. 2:**
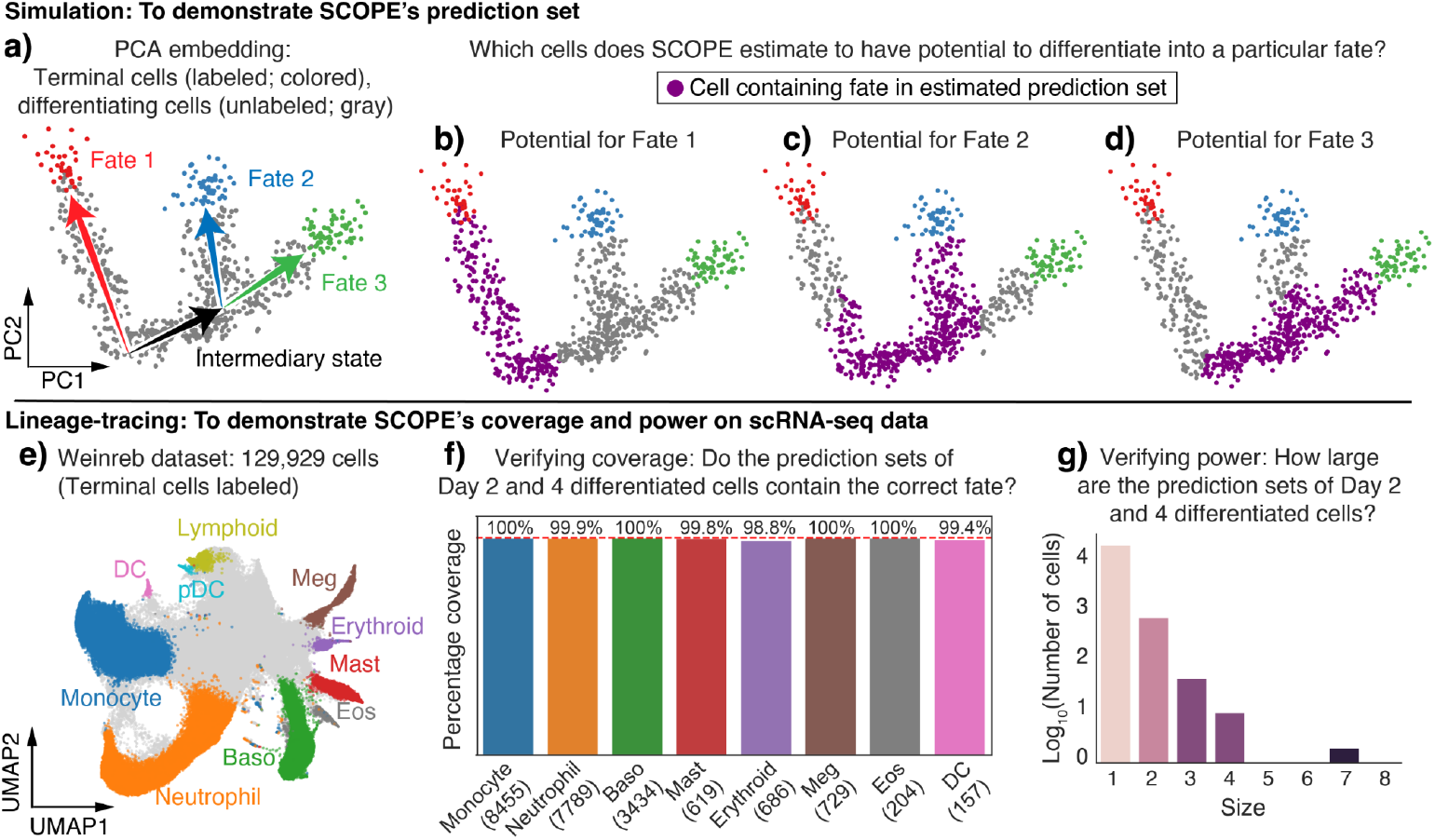
Confirmation of the prediction sets’ coverage and power using simulation and clonal information. **a)** PCA embedding of simulation data, where cells are colored by terminal cell types (red, blue, green) or still undergoing differentiation (gray). **b)** PCA plot showing the cells with Fate 1 in prediction sets. **c)** PCA plot showing the cells with Fate 2 in prediction sets. **d)** PCA plot showing the cells with Fate 3 in prediction sets. **e)** UMAP plot showing terminal cell types of the Weinreb dataset. **f)** Number of Day 2 and Day 4 differentiated cells with their own cell type in their prediction set shown in parentheses, and the percentage coverage, stratified by cell type. **g)** Distribution of the size of prediction sets among all the Day 2 and Day 4 differentiated cells.

We further demonstrated that SCOPE produced robust, well-calibrated prediction sets using clonal and temporal information from a single-cell *in vitro* lineage-tracing dataset of mouse hematopoietic cells derived from bone marrow [20], which we’ll call the “Weinreb dataset.” Heritable clonal (lineage) barcodes of ancestral relationships across different time points provided a natural validation signal, allowing a direct test of whether each cell’s prediction set contained the true fate. In this dataset, 129,929 cells were sequenced across 3 time points (Day 2, 4, and 6), comprising multiple fully differentiated, mature immune cell types as shown in Fig. 2e. We utilized Palantir to generate pseudotime estimates, which will be used to determine recruitment order. The global pseudotime estimation result was shown in Fig. S5. When using SCOPE, we used only Day 6 differentiated cells as the initial training set.

We assessed SCOPE’s prediction sets in two ways. We first checked the coverage of the predictions, answering the question: “Does a cell’s prediction set cover its terminal fate?” Although we used only the differentiated Day 6 cells as labeled cells in SCOPE, the Weinreb dataset also included cells that differentiated at Day 2 or 4. Hence, we expected each cell type that differentiated prior to Day 6 to include its own cell type in its prediction set. Fig. 2e shows the proportion of cells satisfying this criterion by cell type, excluding plasmacytoid dendritic (pDC) and lymphoid cells due to their small numbers. Each terminal cell type had between 8 and 8,455 Day 2 to Day 4 cells (mean: 2,705). For each cell type, at least 99% of cells had prediction sets that included their true label, indicating well-calibrated coverage of the constructed prediction sets.

We next check SCOPE’s power to answer the question, “Does a cell’s prediction set localize to its true fate?” This was equally important to check, since a prediction method that outputted all possible fates would have perfect coverage but be biologically uninformative. To assess power, we investigated the prediction set size of each differentiated cell at Day 2 and Day 4. We expected that a high-powered prediction would have many prediction sets of size 1. Fig. 2f confirmed this, where 96.2% of Day2 and Day4 differentiated cells have a prediction set of size 1. Additionally, we investigated clonal barcodes of multipotent fates. Consider an example clone that has undifferentiated cells at Day 4 and both erythroid (Ery) and megakaryocyte (Meg) cells at Day 6. We expect that the Day 4 prediction set for each undifferentiated cell in this clone should have contained at least one of {Ery, Meg} . We performed this analysis across all clones and found that 96% of Day 4 undifferentiated cells met this criterion. Furthermore, there was a smooth continuum in the number of unitpotent and multipotent cells covering all lineages, reflecting SCOPE’s power to meaningfully recover the complexity of hematopoiesis differentiation (Fig. S6).

### 2.3 SCOPE localizes branchpoints for hematopoiesis

After confirming the statistical validity and power of SCOPE’s prediction sets, we demonstrated how these sets can localize branchpoints in a differentiating system. To do this, we investigated the prediction sets of *all* the cells in the Weinreb dataset, including the undifferentiated cells at Days 2 and 4. These undifferentiated cells represent multipotent cells that differentiate into various cell types in the hematopoietic system (Fig. 3a, [35]), and we were interested in assessing the degree to which each cell is individually specified for a future fate. SCOPE’s prediction set and density estimation, both essential components for branchpoint localization, offered complementary perspectives of this (Fig. 3b,c). From a bird’s eye view of the results in Fig. 3b, we observed that the larger prediction sets are further away from the terminal cells. This was a meaningful diagnostic, since it suggested that SCOPE was appropriately capturing the multipotency of undifferentiated cells the “further” they are from differentiated cells along this cellular manifold. The estimated density demonstrated high-density regions among the undifferentiated cells reinforced our premise that branchpoints occur in high-density regions, especially in hematopoietic systems, where a series of checkpoints tightly control the differentiation of undifferentiated cells [36, 37] (Fig. 3c). However, there were also high-density regions within differentiated populations, indicating that not every high-density region was a branchpoint. The novelty of SCOPE lies in combining prediction sets and high-density regions to rigorously identify branchpoints.

**Fig. 3:**
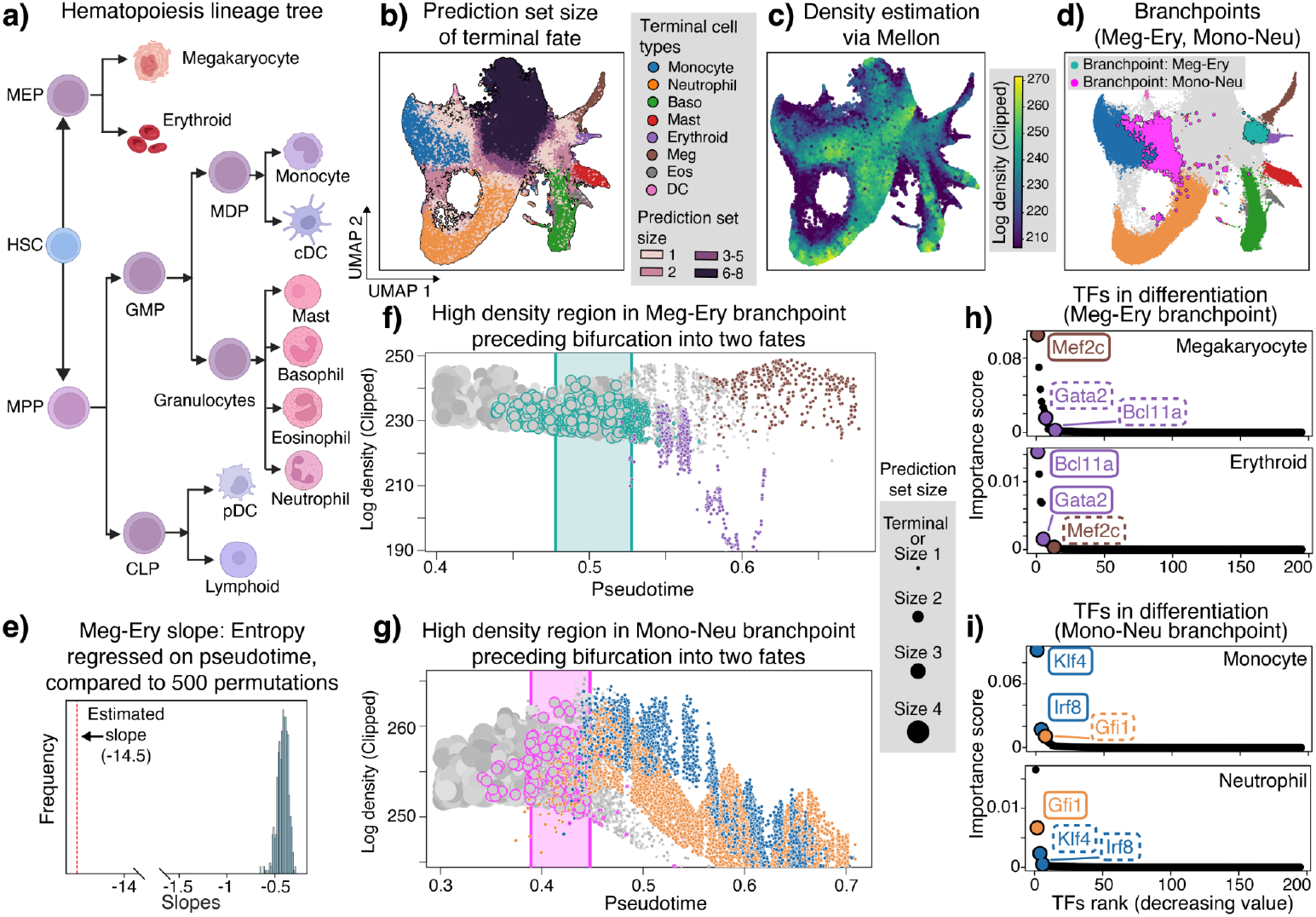
Investigation of branchpoints using lineage-tracing scRNA-seq data. **a)** The lineage tree of hematopoiesis, generated via Biorender (HSC, hematopoietic stem cell; MPP, multipotent progenitor; MEP, megakaryocyte-erythroid progenitor; GMP, granulocyte-monocyte progenitor; CLP, common lymphoid progenitor; MDP, monocyte-dendritic progenitor; cDC, conventional dendritic cell). **b)** UMAP embedding showing cells at the terminal cell type and size of prediction sets for all unlabeled cells. **c)** Cell state density estimation. **d)** Megakaryocyte-erythroid branchpoint and monocyte-neutrophil branchpoints. **e)** Distribution of sampled slopes for the megakaryocyte-erythroid branchpoint when compared to a null distribution. **f)** Cells before and after the branchpoint the megakaryocyte–erythroid branchpoint, plotting the log density against pseudotime for cells. The color and size of each cell depict its cell type and the size of its conformal prediction set. Cells inside the localized region in the branchpoint are highlighted, and the 25th-75th quantile of the branchpoint pseudotimes are depicted as a band for visual aid – not all cells within this band are part of the branchpoint. Only a subset of cells with megakaryocyte or erythroid in their prediction sets are shown for visual clarity. **g)** Cells before and after the monocyte-neutrophil branchpoint, plotted similarly to (f). **h)** Importance scores reflecting how strongly a gene contributes to a cell’s differentiation toward either megakaryocyte or erythroid terminal fate among the cells at the branchpoint. The known driver TFs for each lineage have a bold outline, while driver TFs for the other lineage under consideration in the branchpoint have a dotted outline. **i)** Importance scores across cells at the monocyte-neutrophils branchpoint.

While SCOPE was able to identify branchpoints anywhere along the differentiation tree, we focus on the branchpoints between specific pairs of cell types here. In particular, we analyzed two branchpoints: one between the differentiation between the megakaryocyte and erythroid lineages (cyan), and another between the monocyte and neutrophil lineages (pink; Fig. 3d). While it was expected to see that the localized branchpoints lie “in between” their respective two terminal cell types in the cellular manifold, SCOPE was able to identify the *specific* cells and their associated transcriptomic profile that define the branchpoint. For instance, the branchpoint between monocytes and neutrophils had a clear skew towards the monocyte transcriptomic profiles (Fig. 3d). This highlighted our broad finding that the branchpoints in differentiation are not simply the “midpoint” between the two terminal cell types, as implicitly assumed by other methods such as Slingshot [7] and K-Branches [6]. Analysis of the clonal barcodes of branchpoint cells demonstrated an enrichment of clones that were bipotent for both monocytes and neutrophils, further reinforcing the validity of our estimated branchpoint (Fig. S4).

We next sought to reinforce the premise that SCOPE’s branchpoints represented localized regions where fate specification changed rapidly using an orthogonal evaluation metric and visualizations. Our metric assessed how the entropy of predicted fate probabilities changes within the branchpoint, since a branchpoint biologically represents a large shift from being neutral across multiple fates (high entropy) to specifying a subset of fates (low entropy). For example, the slope when regressing entropy against pseudotime for the megakaryocytes-erythroids branchpoint was -14.5 (Fig. 3e). This was significant when compared with a null distribution of 500 equal-sized subsamples of randomly chosen cells with prediction sets {Meg}, {Ery}, or {Ery, Meg} (p-value = 0.002). Visualizations provided additional context. We plotted the cells’ density along pseudotime, highlighting which cells were localized to the branchpoint in either the Meg-Ery or Mono-Neu branchpoint (Fig. 3f,g). These plots reflected how the branchpoints were defined by a high-density region of bipotent cells preceding fate specification. For example, while high-density cells were still transcriptomically undifferentiated earlier in pseudotime than most megakaryocytes (around pseudotime 0.55), these cells were already specified for the megakaryocyte fate. We further analyzed the branchpoints of other analogous analyses on other branchpoints (Fig. S5).

Lastly, we showcased the biological relevance of localized branchpoints by interrogating transcription factors (TFs) established as master regulators of hematopoiesis. Using transcript abundance serves as an operational proxy for protein activity, we investigated if these TFs driving fate specification had a high importance score in SCOPE’s respective fate classifier. Indeed, at the Meg-Ery branchpoint, SCOPE correctly prioritized *Mef2c* in the megakaryocyte classifier [38], while *Gata2* and *Bcl11a* [39–41] showed higher importance for the erythroid fate, which recapitulates known drivers of these fates (Fig. 3h). This pattern of fate-specific reversal was also displayed at the Mono-Neu branchpoint, where monocyte drivers *Klf4* and *Irf8* [42, 43] were prioritized over the neutrophil driver *Gfi1* [44] in their respective fate classifiers (Fig. 3i). This reversal in importance scores demonstrates that SCOPE accurately resolves the distinct regulatory programs that define lineage specification.

We additionally compared SCOPE against Palantir (Fig. S8), CoSpar ([45]; Fig. S10 and S9), and the entropy of CellRank ([9]; Fig. S11) based on their ability to localize branchpoints. Although CoSpar and CellRank leverage biological information not available to SCOPE, we observed that these methods still struggled to localize the branchpoint to a specific transcriptomic region.

### 2.4 Biological replicates demonstrating the consistency of the branchpoints

We next demonstrated that SCOPE infers reproducible branchpoints by inferring consistent biological pathways and differentially expressed genes (DEGs) across biological replicates. We analyzed two CD34^+^ human bone-marrow datasets from health donors [8, 21], which we refer to as the “Setty” and “Persad datasets” (4,142 and 6,881 cells of erythroid, monocyte, and dendritic cells, respectively), and focused on the specification between the erythroid and monocyte fates. To facilitate comparisons between the two datasets, we sought to confirm the consistency of SCOPE by nominating the genes that were differentially expressed between the two fates within the branchpoint (Fig. 4a).

**Fig. 4:**
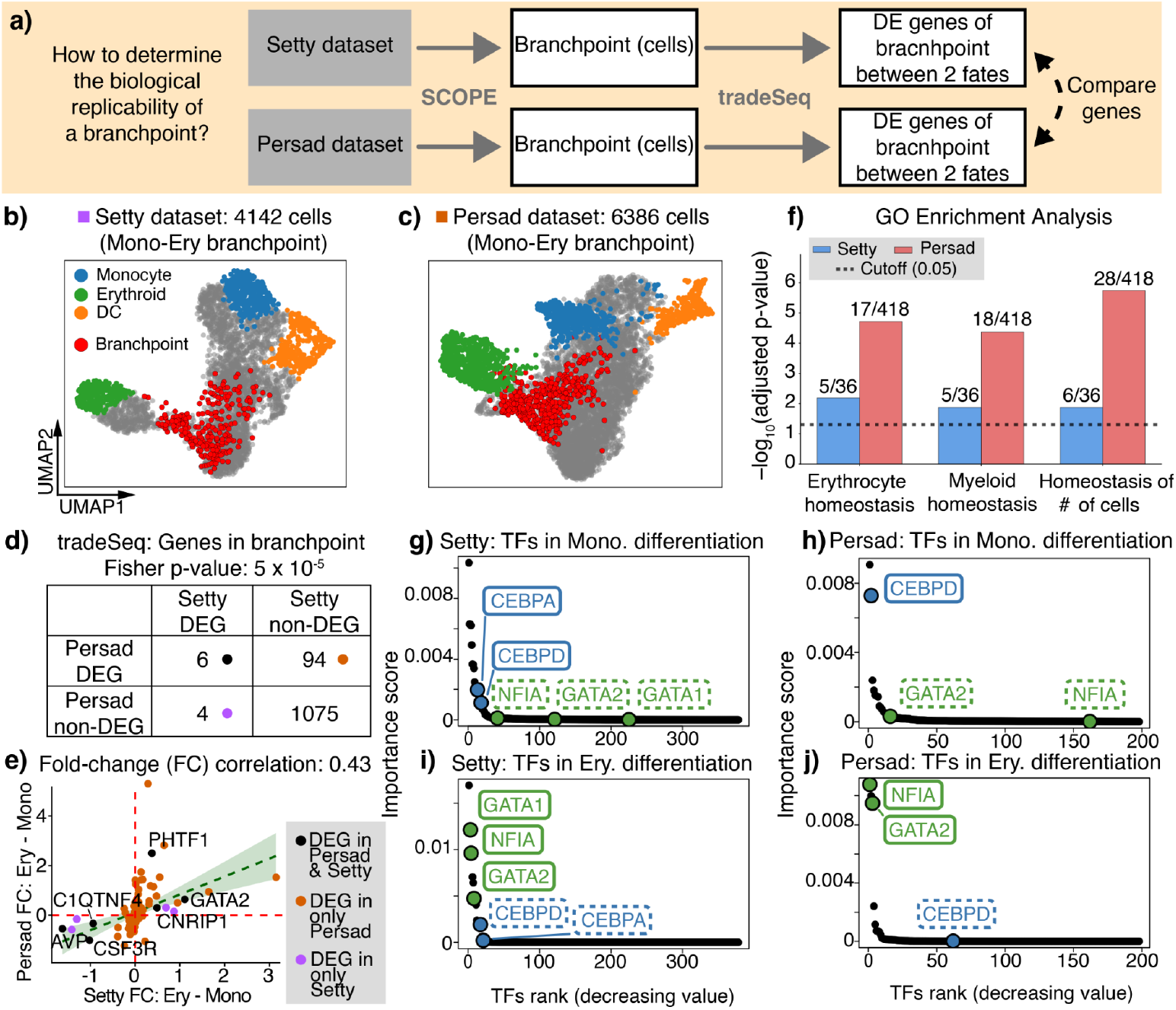
Biological replicability of SCOPE’s branchpoints across different hematopoiesis datasets. **a)** Workflow of validating SCOPE’s branchpoints. **b**,**c)** Branchpoint between erythroid and monocyte for the Setty and Persad dataset, respectively. **d)** Comparison of tradeSeq’s differentially expressed genes between the two lineages at branchpoints. **e)** Correlation of fold changes between the erythroid and monocyte lineages across the two datasets, among genes found in at least one dataset. **f)** All GO enriched pathways found in both datasets, where the fraction denotes how many DEGs in the pathway were identified among the total pathway size. **g**,**h)** Importance scores for the monocyte fate within the branchpoint for Setty or Persad, respectively, where only human TFs are shown. Key TFs for granulocyte-monocyte (blue) or erythroid (green) differentiation are colored, with a plotting similar to Fig. 3h,i. **i**,**j)** Same format as (g,h), but for the erythroid fate.

We first used SCOPE to localize the branchpoint in each dataset (Fig. 4b,c). We then applied tradeSeq [46] to identify differentially expressed genes (DEGs) between the two lineages within the branchpoints. Briefly, tradeSeq fitted regression models along developmental trajectories to identify genes that were differentially expressed between erythroid and monocyte lineage specification, weighting each cell by its fate bias. When we compared the implicated DEGs between the two datasets, we found a significant concordance (Fig. 4d; Fisher’s exact test p-value = 5.0 × 10^*−*5^). Additionally, for the 104 DEGs implicated in either dataset, we calculated the fold change between the two fate biases (see Supplementary Note S6). The fold changes were significantly positively correlated (Pearson correlation = 0.43) between the two datasets (Fig. 4e). *GATA2*, a gene implicated in both datasets, had a strong erythroid bias, reinforcing its role as a TF critical for erythropoiesis [39, 47]. We next wondered whether the implicated DEGs encoded proteins involved in similar biological processes. We performed a GO enrichment analysis based on the dataset-specific DEGs for each dataset to further confirm key biological pathways that were significantly enriched in both (Fig. 4c). All three significant pathways overlapping in both datasets (erythrocyte homeostasis, myeloid cell homeostasis, and homeostasis of cell number) were biologically related to the branchpoint between the erythroid and monocyte lineages. These pathways were collectively driven by the genes *GATA2, KLF1, RPS19*, and *B2M* in both datasets. These findings collectively confirmed the consistency of the monocyte-erythroid branchpoint across independent datasets at the gene level.

Having assessed the branchpoint similarity between the two datasets, we next investigated the relative importance of known transcription factors critical for the monocyte or erythroid differentiation within our branchpoints. To do this, we applied our importance score metric described in Section 2.3 and ranked only the genes associated with human TFs [48]. Based on a broad literature search, among the genes highly variable in our dataset, we labeled GATA1, GATA2, and NFIA as the drivers of erythroid differentiation [39, 40, 47, 49, 50], and CEBPA and CEBPB to a lesser extent as the drivers of granulocyte-monocyte differentiation [51, 52] (see Fig. 3a). First, when investigating SCOPE’s classifier for the monocyte fate, we observed that *CEBPA* and *CEBPD* had a higher importance than *GATA1, GATA2*, and *NFIA* (Fig. 4g,h). We note that *CEBPA* and *GATA1* were not highly variable genes in the Persad dataset. Second, when investigating SCOPE’s classifier for the erythroid fate, we observed a reversal in importance (Fig. 4i,j). Importantly, in many of our comparisons, our labeled TFs were among the first or second most important TF. This replicable inversion of transcription factor importance across both datasets showcased SCOPE’s ability to correctly resolve fate-specific regulatory programs.

Additional results were shown in Fig. S13 and S18, where we performed a similar analysis described in Section 2.3 to show that branchpoints represented groups of cells dynamically undergoing fate specification. Furthermore, we demonstrated in Fig. S15 to S16 how Slingshot struggles to discover robust branchpoints on the Setty dataset.

### 2.5 SCOPE uncovers epigenetic priming in hematopoiesis and retina

While branchpoint localization identified the cellular states immediately preceding specification, characterizing the molecular logic of these transitions required resolving when regulatory signals first become fate-informative. We focused on *epigenetic priming*, a state of regulatory poise where chromatin accessibility at enhancers or promoters foreshadows the transcriptional execution of a lineage program [1]. Although characterizing this temporal lag is a central goal of multi-omic workflows [10, 21], current methods typically quantify priming by identifying asynchronous changes in raw expression levels – a metric that may not directly link molecular shifts to a specific fate decision. SCOPE addressed this by using the dynamics of calibrated prediction sets as a robust, informational metric of timing: if chromatin signals drive specification, the ATAC-trained classifiers should resolve fate resolution (e.g., singleton prediction sets) earlier in pseudotime than their RNA counterpart. This set-based comparison provided a broad overview of modality lag, but the framework’s unique utility resided in its additional ability to localize the specific regulatory drivers underlying these timing shifts (Fig. 5a). By interrogating the importance scores of transcription factors and their linked enhancers that drive the underlying prediction, SCOPE prioritized a feature’s informational contribution to a fate decision rather than its raw expression level. This offered a fate-specific lens that complements existing biophysical models.

**Fig. 5:**
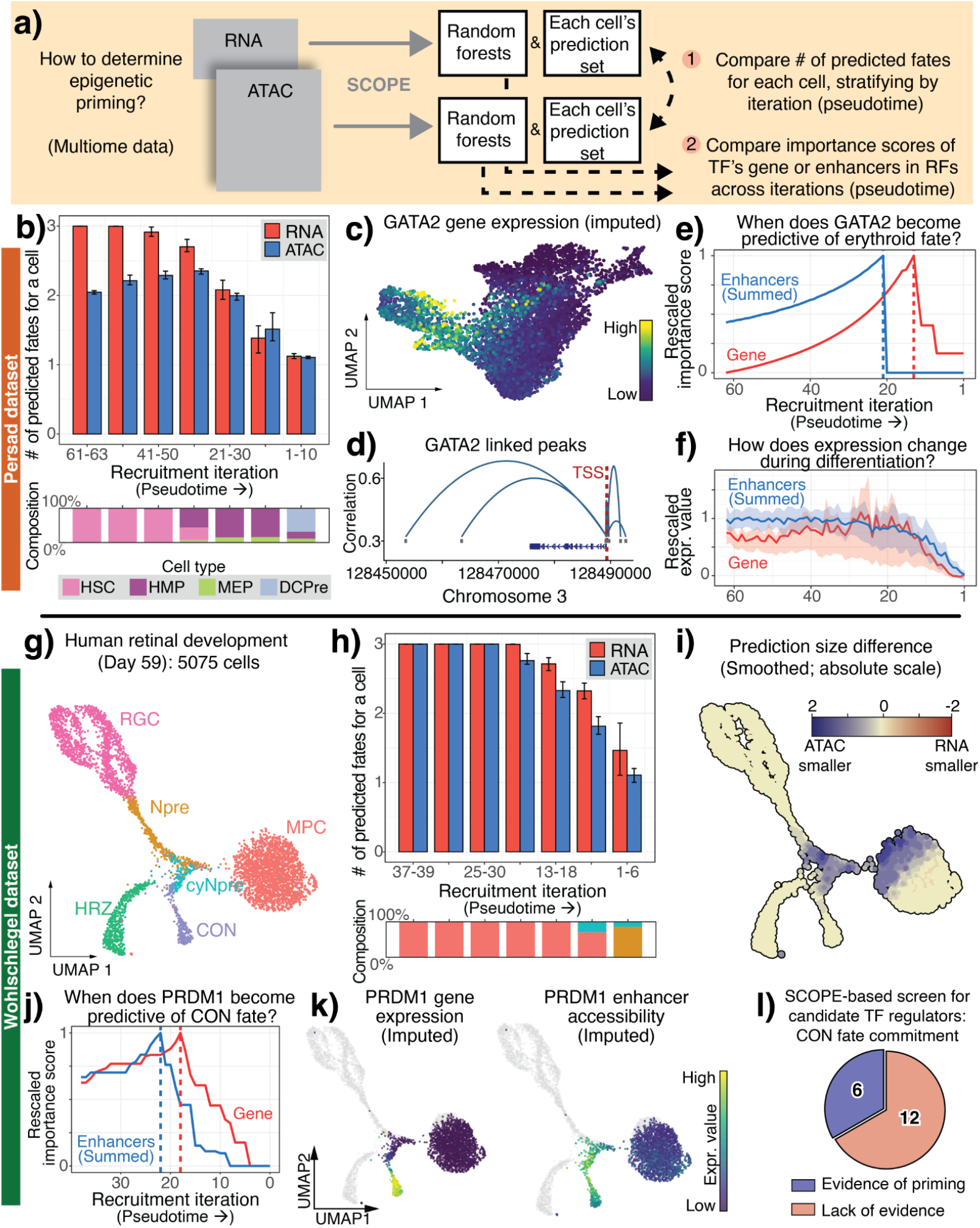
Assessment of epigenetic priming via SCOPE of erythroid differentiation and cone photoreceptor differentiation. **a)** Workflow of deriving priming, both genome-wide (via prediction set size) and TF-specific (via importance scores). **b** Prediction set sizes for cells across recruitment iterations with 25th-75th quantile bands, shown with the cell-type composition within each bin of recruitment. **c**,**d)** GATA2’s gene expression, a driver of erythroid differentiation, and its linked peaks learned from the Persad dataset (DCPre, dendritic cell precursor). **e**,**f)** GATA2’s importance scores or rescaled expression values for both gene expression and enhancers based on the erythroid classifier across recruitment iterations, respectively. The median normalized expression is shown in (d), rescaled so that the trend lies in [0, 1], with the 25th-75th quantile bands overlaid. **g)** UMAP of Wohlschlegel dataset (MPC, multipotent progenitor cells; Npre, neurogenic precursors; cyNpre, cycling neurogenic precursors; HRZ, horizontal cells; CON, cone photoreceptors; RGC, retinal ganglion cells). **h)** Same plotting layout as (c) but for the Wohlschlegel dataset. **i)** Smoothed difference in prediction set sizes (RNA-ATAC) visualized on the UMAP. **j)** PRDM1’s importance scores for both gene expression and enhancers based on the cone photoreceptors’ classifier across recruitment iterations. **k)** UMAPs showing the imputed PRDM1 gene expression or summed enhancer accessibility, only among cells with cone photoreceptors in its prediction set. **l)** Proportion of 18 TFs for the cone photoreceptor fate that display epigenetic priming among TFs that were highly variable and had detectable linked enhancer regions.

To evaluate SCOPE’s capacity to detect broad epigenetic leads, we returned to the multi-omic Persad bone-marrow dataset. After separately applying SCOPE to the ATAC modality (where features corresponded to peaks denoting open chromatin regions) and the RNA modality, we observed a systematic difference in the timing of fate specification. Across the three terminal fates, ATAC-derived prediction sets among hematopoietic stem cells (HSCs) resolved toward singletons substantially earlier in pseudotime than their RNA counterparts (Fig. 5b). This broad cell-level lead suggested that chromatin signals provided a signature of specification earlier during differentiation than the transcriptome. To investigate the molecular drivers of this lead, we focused on GATA2, which SCOPE identified as a top-ranked erythroid driver in both bone-marrow replicates (Fig. 4i,j). While *GATA2* expression remained low until cells reach a differentiated erythroid state (Fig. 4c, 5c), we found that the importance of its linked cis-enhancer peaks (Fig. 5d) was consistently high in HSCs and hematopoietic multipotent progenitors (HMPs), reaching its informational apex significantly earlier in pseudotime than the importance of the gene’s expression (Fig. 5e). Notably, this priming evidence was largely masked when examining imputed expression levels alone, which remained relatively flat across early progenitor states (Fig. 5f). These results demonstrated that SCOPE identified regulatory poise by leveraging fate-specific importance scores that account for the collective information contributed by all enhancers simultaneously, rather than studying each set of enhancers in isolation. See Supplementary Note S4 for more details on preprocessing.

To demonstrate that SCOPE was generalizable across biological systems with distinct developmental dynamics, we next investigated human retina development using the multi-omic dataset (fetal day 59, 5,075 cells; Fig. 5g) [22, 53], which will refer to as the “Wohlschlegel dataset.” Consistent with our findings in human hematopoiesis, the retina system exhibited clear evidence of epigenetic priming, with ATAC-derived prediction sets resolving toward terminal fates significantly earlier than their RNA counterparts (Fig. 5h,i; see Fig. S20 for the specific prediction sets). However, we observed a notable shift in global timing: while the modality lag in the bone marrow manifested in the earliest progenitor states, the largest differences in prediction set resolution in the retina occurred later in pseudotime. To determine whether this global lag was driven by a similar regulatory logic observed in hematopoiesis, we interrogated PRDM1, a known transcription factor driver of cone photoreceptor (CON) differentiation [54]. Contrasting the two modalities’ importance trajectories revealed that the informational apex of PRDM1 enhancers occurred before the apex of the gene’s transcriptomic importance (Fig. 5j. We note that while these importance scores effectively localized the predictive power and timing of regulatory elements, they are agnostic to the direction of the biological effect; rather, it was the subsequent integration with imputed expression values (Fig. 5k) that identified PRDM1 as an activating driver of the cone photoreceptor lineage. This temporal offset reinforced the notion that SCOPE reliably localized the molecular drivers of fate specification in diverse biological systems.

Beyond interrogating individual drivers, SCOPE’s importance framework enabled a systematic, genome-wide screen for potential epigenetic priming signals. To establish this utility, we conducted an automated survey of 18 transcription factors driving cone photoreceptor differentiation, classifying their timing based on the relative timing of their informational apexes. These were all TFs with highly variable genes and at least one significantly linked peak enhancer. Using a lead of at least one recruitment iteration as a criterion, we identified six high-priority TFs exhibiting clear evidence of epigenetic priming, including PRDM1, DLX1, and NR2E1 (also known as TLX) (Fig. 5l) [55]. These results provided a prioritized list for functional follow-up validation. To contrast this list, other key TFs during retinogenesis not specific to the cone photoreceptor differentiation, such as NEURDO1 and ONECUT2, remained ambiguous in our analysis [56–58]. Interestingly, although we observed a global lead for the ATAC modality at the cell level (Fig. 5h), our screen suggested this predictive power was anchored by a selective minority of TFs rather than a uniform transcriptomic shift.

## 3 Discussion

SCOPE establishes a statistically rigorous framework for localizing developmental branchpoints and regulatory priming by transforming fate probabilities into calibrated prediction sets. By formalizing “poise” through conformal inference, SCOPE identifies commitment checkpoints where fate-ambiguity and high cell-state density co-occur. However, our approach has some limitations. Our density-based localization reflects a convolution of differentiation “speed” and sampling frequency, requiring careful interpretation in asynchronous systems since density peaks may not necessarily perfectly align with decision coordinates. It also requires pre-specified terminal fates, since all of SCOPE’s results are relative to the labeled terminal fates of interest. Furthermore, while SCOPE’s multi-omic versatility enables the detection of epigenetic priming, the precision of these temporal lags is inherently limited by the resolution of peak-calling and the biological stage of development represented in the multi-omic dataset. For example, our TF screen in Fig. 5l was conducted exclusively on cells from fetal day 59, so TFs that did not exhibit priming may suggest potential regulatory lags that extend beyond our specific temporal snapshot. Future work could integrate SCOPE’s framework with a regulatory network inferred from paired multi-omic data to strengthen enhancer signals, eliminating the need to aggregate enhancers to obtain a robust importance score.

Despite these limitations, SCOPE’s modality-agnostic nature allows it to serve as a foundation for integrating spatial and chromatin contexts into a unified developmental map. By replacing heuristic thresholds with distribution-free coverage guarantees, SCOPE provides the resolution necessary to identify developmental “poise” in systems beyond those analyzed in this paper, such as human organoid morphogenesis, peripheral immune-cell exhaustion, and state changes in microglia during neurodegeneration. Ultimately, this framework moves single-cell analysis beyond descriptive trajectories toward a predictive, coordinate-based understanding of the molecular logic governing cell-fate commitment.

## Method and analysis

The following is an overview of SCOPE that explains the key details of each item in the subsequent sections. See the pseudocode in Supplementary Note S5..

1. **Preprocessing**: SCOPE requires three key inputs, which are detailed more in Supplementary Note S4 for the specific datasets:
  a. **Normalized single-cell feature matrix**: Let *X* ∈ ℝ^*n×p*^ be a dataset with *n* cells and *p* highly variable genes, where *X*_*ij*_ denotes the denoised library-size-adjusted gene expression of gene *j* in cell *i*. Broadly speaking, we use `Scanpy` [59] to determine the highly variable genes, and use scVI [60] or peakVI [61] to normalize the count matrix of RNA or ATAC, respectively, whenever possible. If the count matrix is unavailable, a log-normalized matrix can be used as input.
  b. **Pseudotime estimate for each cell**: SCOPE requires pseudotime to determine cell recruitment order, and we use Palantir [8] as the default method. This method takes the top principal components of the RNA modality’s normalized gene expression matrix and learns a temporal ordering using a diffusion map. Let *t*_*i*_ ∈ [0, 1] denote the estimated pseudotime of cell *i*.
  c. **Annotation of terminal cell types**: SCOPE requires knowledge of what the terminal cells are in a particular single-cell dataset. For datasets without such annotations, we perform Louvain clustering using high resolutions and assign terminal cell types based on known marker genes. Let *m* be the number of terminal cell types.
2. **Iterative recruitment**: SCOPE quantifies the differentiation potential of unlabeled cells by iteratively recruiting cells in pseudotime order (from most terminal-like to most progenitor-like) and generating prediction sets for each unlabeled cell. Starting cells at high pseudotime, we recruit a subset of unlabeled cells with an earlier pseudotime. Hence, in each iteration, we have labeled cells (which we split into a “training set” and a “calibration set”) and unlabeled cells (of which a subset will become the “test set”, i.e., cells we recruited into this iteration). After we form the prediction set of all cells in the test set, those cells become part of the labeled cells for the next iteration.
3. **Multi-label binary random forest**: To perform these predictions, SCOPE employs a multi-label binary-relevance random forest framework that can model prediction sets containing multiple cell types. See Supplementary Note S3 for a more in-depth discussion about the advantage of using a random-forest classifier over a deep-learning classifier for SCOPE. Specifically, this ensemble trains one binary classifier per terminal cell type on the training set, where each classifier learns to determine whether a given cell’s prediction set includes that fate. The models are trained iteratively with a warm start, which adds new trees at each iteration while retaining previously learned parameters. For each unlabeled cell, the trained classifiers output probabilities across all terminal types, which are then normalized to yield a probability distribution over cell fates.
4. **Label propagation**: After predicting at each iteration, we smooth the predictions using an adjacency graph built from all cells [24], a common approach in semi-supervised learning. This propagation is key to SCOPE and enables the labels of the training cells to “spread” along the geometric cell-cell manifold. Without this step, the multi-label binary random forests would incorrectly predict a cell’s fate solely based on transcriptomic similarity between an unlabeled cell and the labeled cells, disregarding the cellular manifold.
5. **Conformal inference and iterative recruitment**: We transform the smoothed fate probabilities into calibrated prediction sets for each cell in the test set using distribution-free conformal inference, yielding multi-fate sets with theoretical coverage guarantees. We use entropy balancing to determine the appropriate weighting of the calibration set for mitigating distribution shift.

Across different iterations, SCOPE recruits unlabeled cells, updates the random forest classifiers (based on the training set) and the calibration scores (based on the calibration set) to form prediction sets on the test set, and repeats this procedure until all cells along the trajectory have prediction sets.

### Iterative recruitment

In our work, we gradually augment the “training set” with originally unlabeled cells that are labeled using the prediction set. Specifically, we bin all the unlabeled cells (i.e., all cells that are not in the terminal cell type) into *L* bins ordered by pseudotime, each of the same size and consisting of cells with consecutive pseudotimes {*t*_*i*_}’s. By default, we set *L* = round 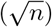 . For a particular iteration, we:

1. Recruit the next bin of unlabeled cells to be the test set. Let this set be denoted as *Te* ⊂ {1, …, *n*}, with size *k*.
2. Construct a calibration set *C* ⊂ {1, …, *n*} of size *c* that is twice the size of the test set from the labeled cells closest in pseudotime to the test set. Let the training set *Tr* ⊂ {1, …, *n*} of size *t* denote all the remaining labeled cells in this iteration that aren’t in the calibration set.

(If there are fewer than 2*k* labeled cells, which might occur in the first few iterations, we set *C* to be half of the labeled cells.)

### Multi-label binary random forest

Since prediction sets may contain multiple cell types (non-singleton sets), we need a multi-label classification framework. Given *m* terminal fates, we train *m* independent binary classifiers using random forests. Empirically, we have found that training *m* separate random forest classifiers performs better than other classification strategies for modeling multi-label outcomes where the outcome is a set (i.e., possibly more than one predicted label), as discussed in Supplementary Note S3.

During training, for each terminal fate *j* ∈ {1, 2, …, *m*}, we train a binary random forest classifier *f*_*j*_ on training set where:

- Positive examples: cells whose prediction set contains terminal fate *j*.
- Negative examples: cells whose prediction set does not contain terminal fate *j*.

During testing, for an unlabeled cell, each trained classifier *f*_*j*_ for terminal fates *j* ∈ {1, …, *m*} outputs a probability *p*_*j*_ ∈ [0, 1] representing the likelihood that the cell differentiates into terminal fate *j*. The normalized probability distribution over the *m* fates is:

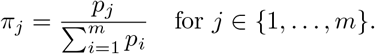

We use the `RandomForestClassifier()` function from the `scikit-learn` package to train our random forests. See Supplementary Note S6 for more details.

### Label propagation

Due to the distribution shift, the learned prediction patterns from the training set are not directly transferable to the calibration and test sets. This results in poor predictive performance and motivates the need for a semi-supervised approach. At a high level, label propagation “smooths” the predicted probabilities from the multi-label binary random forest classifier to conform to the cell-cell manifold (i.e., a graph connecting cells with similar transcriptomic profiles). This provides probability estimates for the adaptive conformal scores that are less affected by the distribution shift. This label propagation consists of two steps: first, construct the graph; second, iteratively smooth the predictions along it after each iteration.

#### Step 1: Graph construction and normalization

To represent cellular similarity within the high-dimensional manifold, we construct a globally connected graph using a *k*-nearest neighbor (kNN) adjacency matrix using an adaptive Gaussian kernel [8]. The resulting affinity matrix *W* is symmetrically normalized [62] to produce a normalized Laplacian *S*. Detailed mathematical formulations of the graph construction and normalization procedure are provided in Supplementary Note S6.

#### Step 2: Label propagation

Equipped with the cell-cell-normalized adjacency matrix *S*, we propagate information from labeled to unlabeled cells. For a particular iteration, let *Y* ∈ ℝ^*n×m*^ denote the per-cell soft label distributions (i.e., a cell *i* with prediction set 𝒞 (*x*_*i*_), we assign a value of 1*/*|𝒞 (*x*_*i*_)| to all terminal cell types *c* ∈ 𝒞 (*x*_*i*_), and 0 otherwise), ordered as

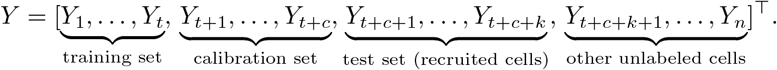

Specifically, the first *t* + *c* rows [*Y*_1_, …, *Y*_*t*+*c*_]^⊤^ are the soft labels of all previously labeled cells (i.e., training and calibration set). This includes the original labeled terminal cells, whose corresponding vector *Y*_*i*_ is a one-hot encoding, as well as all cells recruited in the previous iteration. The remaining rows corresponding to currently unlabeled cells have all values initialized to zero. We then make a copy of *Y* as *Y* ^∗^, the first *t* rows are the same with *Y* . Rows *t* + 1 to *t* + *c* + *k* of *Y* ^∗^ are then updated to be the soft predictions produced in this iteration by a multi-label random forest classifier for the *c* cells in the calibration set and the *k* newly recruited cells (i.e., the test set). The remaining rows are still zero, i.e., 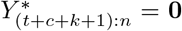. Then we initialize *F* ^(0)^ = *Y* ^∗^ and update

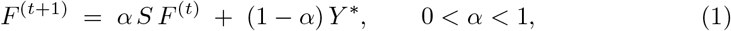

for many iterations until convergence. By default, *α* is set as 0.99. As shown by Zhou et al. [24], the iteration (1) converges to the unique fixed point

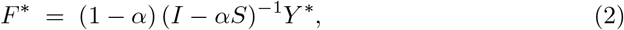

where *I* is the *n* × *n* identity matrix, so in our implementation, we directly use this fixed point solution. Afterwards, we take the rows of 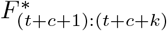 as the final fate probabilities vectors for the newly recruited cells. These fate probabilities are used for the next step (conformal inference). For the next iteration, we update *Y*_(*t*+*c*+1):(*t*+*c*+*k*)_ according to the results of conformal inference for the newly recruited cells.

### Conformal inference and iterative recruitment

To provide a statistically rigorous measure of developmental poise, SCOPE utilizes conformal inference [12, 63] to transform raw fate probabilities into calibrated prediction sets. Unlike standard classification heuristics, this distribution-free framework provides a guarantee that the true terminal fate of a cell is contained within the predicted set with a user-defined probability (1 − *α*). By formalizing fate uncertainty as a set-membership problem, we can define branchpoints as regions where prediction sets remain multi-label (e.g., {Ery, Meg}) and identify epigenetic priming as the regime where these sets resolve toward a single fate earlier in one modality than another. While conformal inference typically provides marginal coverage guarantees, this global calibration is sufficient for SCOPE to reliably localize high-density regions of fate-ambiguity across the differentiation manifold. A formal description of the conformal framework and a discussion of its coverage properties in the context of iterative recruitment are provided in Supplementary Note S1.

#### Adaptive conformal score

Let *y* ⊆ {1, …, *m*} be the (possibly multi-valued) prediction set of a particular unlabeled cell. Suppose the classifier outputs probabilities {*π*_1_, …, *π*_*m*_} (all non-negative and 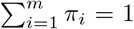 by definition) for this unlabeled cell, and let *π*(1) ≥ · · · ≥ *π*(*m*) be the decreasing rearrangement. Equivalently, let *σ* be the permutation that orders the fates so that *π*(*j*) = *π*_*σ*(*j*)_. Define

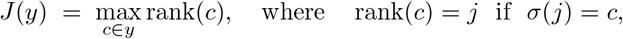

e.g., *J* (*y*) is the worst (largest) rank among the true labels. The conformal score for (*x, y*) is then defined as

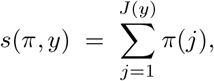

the cumulative probability mass required to cover all true labels. From the calibration set, we obtain scores {*s*_1_, …, *s*_*c*_} and set the threshold to the *α* quantile,

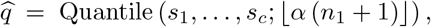

using order-statistic indexing. Note that 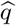 is positive-oriented, i.e. larger values indicate better conformity [64], so we use *α* quantile instead of 1 − *α* quantile.

#### Prediction set for each unlabeled cell

For a test cell with probabilities {*π*_1_, …, *π*_*m*_} and sorted values *π*(1) ≥ · · · ≥ *π*(*m*), define

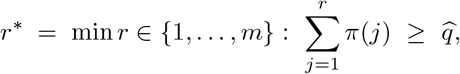

where |𝒜| denotes the size of set . The conformal prediction set is the top-*r*^∗^ fates by probability,

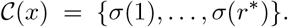

In the idealized setting where calibration set and test set are exchangeable, this conformal construction guarantees that the true fate is contained in the prediction set with probability at least 1 − *α* on average across cells in the test set. As we mentioned in “Step 2: Label propagation”, after constructing the prediction set, we assign a uniform soft label over its elements: each class (i.e., fate) *c* ∈ 𝒞 (*x*) receives probability 1*/r*^∗^, and all classes *c* ∉ 𝒞 (*x*) receive probability 0.

#### Entropy balancing to handle distribution shift

Recruiting cells along pseudotime {*t*_*i*_} ‘s induces a (transcriptomic) distribution shift between the calibration and test sets, so the exchangeability assumption required by standard conformal inference does not hold. To mitigate this, we reweight the calibration cells using entropy balancing [25, 65] so that key features of the calibration set match those of the test set. This method is commonly used in causal inference to account for distribution shifts.

Formally, suppose the calibration set *C* contains *c* cells with gene-expression profiles 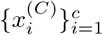 and the test set *Te* contains *k* cells 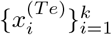, where *x* ∈ ℝ^*p*^. The goal is to compute nonnegative weights *w* = (*w*_1_, …, *w*_*c*_) on calibration cells with 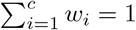 that enable the weighted calibrated cells to have a similar quality to the test cells. Let *f* : ℝ^*p*^ → ℝ^*m*^ denote a feature map capturing quantities to be balanced across sets. In our application, *f* (*x*_*i*_) ∈ ℝ is the Shannon entropy of the predicted fate probabilities for cell *i*, quantifying uncertainty in fate specification (i.e., *p* = 1). We then choose maximum-entropy weights subject to feature matching:

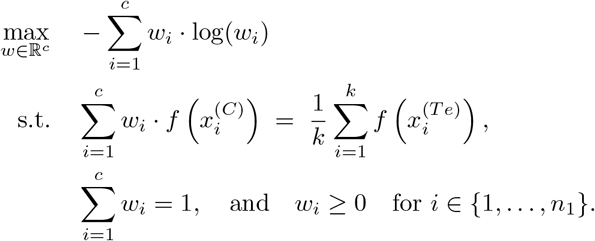

The resulting weights are used to define the weighted quantile 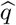 during conformal inference, which realigns the calibration-set feature distribution with that of the test set while avoiding degenerate solutions that place all probability mass on a single cell. We solve the above optimization problem using CVXP [66].

### Downstream: Localization of branchpoints via density clustering

#### Estimation of density in transcriptomic space via Mellon

Mellon [26] is a method for estimating cell-state densities from high-dimensional single-cell data, enabling the identification of dense, stable cell types and sparse, transitional states that connect them. We apply Mellon to estimate per-cell density for all our datasets in this paper, using the `DensityEstimator()` and `fit_predict()` functions from the `mellon` package. See Supplementary Note S6 for more details.

#### Density clustering

The idea of density clustering here is basically the same as [67]. To find a branchpoint between Fate *A* and Fate *B*, we first subset all cells with prediction set {*A*}, {*B*}, {*A, B*} . Then, we cluster the cells based on the estimated density via an iterative hillclimbing method. Broadly speaking, cells “flow” to neighboring cells with a higher density until they reach the cell with the largest local density (i.e., cluster center). The clusters are then defined by the basins of attraction of their respective cluster centers. We modify this procedure to account for the conformal prediction sets. Then, for each such cell, we connect it to its *k* nearest neighbors (default *k* = 30) that satisfy both: (i) higher density; (ii) a prediction set that is equal to or a superset of the cell’s own set. A cell with no admissible neighbor other than itself is designated a cluster center. The branchpoint will be the cluster with the largest number of cells with the prediction set of {*A, B*}.

#### Beyond branchpoints between two terminal fates

We note that in our primary analyses, we focused on two fates, *A* and *B*, which were specific cell types. Conceptually, SCOPE can treat *A* and *B* as multi-labels of terminal fates (for example, *A* being any cell predicted to differentiate to specifically {Monocyte, Neutrophil}, and *B* being any cell predicted to differentiate to only {Basophil} cells, respectively). This type of “inner” branchpoint is similarly analyzed in other methods such as Carta [16]. We perform an analysis of these inner branchpoints in Fig. S7.

### Downstream: Differential expressed genes within a branchpoint via tradeSeq

After localizing branchpoints between two terminal fates, we aim to identify the driver genes responsible for the divergence of cell fates. To achieve this, we use the `tradeSeq` package, [46] to fit regression models along developmental trajectories and identify genes that are differentially expressed along one or multiple fates. Specifically, we use the `fitGAM()` function to detect genes that are differentially expressed between the two fates within each branchpoint. This procedure requires pseudotimes and cell-specific weights, which we use as the pseudotime estimates {*t*_*i*_} ‘s from Palantir and its estimated fate bias, respectively. Then, we use the `patternTest()` function to test for genes that are statistically different between the two fates. The resulting *p*-values are adjusted for multiple testing using the Benjamini-Hochberg (BH) procedure. See Supplementary Note S6 for details.

### Downstream: Importance scores to assess features informative for a branchpoint

Since SCOPE iteratively builds random forest–based classification models to predict the fate of unlabeled cells, our framework naturally enables quantification of gene importance across the iterations that encompass the cells within a given branchpoint. Recall that our classifier consists of *m* binary random forests, one for each fate, which allows us to compute fate-specific feature importance scores. Suppose the branchpoint consists of cells across *k* different recruitment iterations (see “Iterative recruitment”). We denote *n* = (*n*_1_, · · ·, *n*_*k*_) as the number of cells in each iteration across branch-point, and 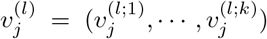 as the importance scores for gene *j* from the classifier corresponding to fate *l* across *k* iterations. Importance scores are obtained using the `feature_importances_` attribute from the `scikit-learn`’s random forest. These scores are calculated as the Mean Decrease Impurity, which quantifies the total reduction in node impurity attributed to a specific feature [68, 69]. The normalized importance of gene *j* for fate *l* is defined as the weighted average of importance scores across iterations, where the weights are calculated as the proportion of cells in each iteration:

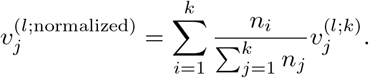

### Downstream: Assessment of presence/absence of epigenetic priming

We document our usage of linking enhancers to genes via SEACells [21] in Supplementary Note S6. After linking enhancers to genes, we assess whether the chromatin is poised (i.e., primed) by repurposing the importance score we designed in the “Importance scores to assess features informative for a branchpoint” section.

Specifically, we first linearly rescale the importance score of a feature (i.e., gene or enhancer) to the range [0, 1] since our assessment of epigenetic priming is based on the recruitment iteration at which the importance score reaches its apex. When analyzing the enhancer regions of a TF, we summed the importance scores across all linked enhancer regions prior to linear rescaling. Then, we fit a unimodal regression model using `ufit()` from the `Iso` package [70] with `type = “raw”` to the normalized variable-importance trajectories (rescaled to have values between 0 and 1) of a TF’s gene expression and their corresponding enhancer accessibility across iterations across pseudotime. This is to smooth the importance score and have a clearly defined apex. This unimodal regression is fit separately for the gene expression and the enhancer feature. Epigenetic priming is defined as cases in which the mode of the enhancer trajectory occurred more than one iteration earlier than the mode of the corresponding TF trajectory.

## Data availability

The Weinreb dataset [20] is available at https://github.com/AllonKleinLab/paper-data/tree/master/Lineage_tracing_on_transcriptional_landscapes_links_state_to_fate_during_differentiation. The Setty dataset [8] is available at https://dp-lab-data-public.s3.amazonaws.com/palantir/marrow_sample_scseq_counts.h5ad. The Persad dataset [21] is available at https://dp-lab-data-public.s3.amazonaws.com/SEACells-multiome/cd34_multiome_rna.h5ad. The Wohlschlegel dataset [22] is available at https://www.ncbi.nlm.nih.gov/geo/query/acc.cgi?acc=GSE246169.

## Code availability

The code of SCOPE and the analyses to reproduce our results are publicy available at https://github.com/YiminZhao97/SCOPE.

## Acknowledgements

We thank Ali Shojaie, Wei Sun, Michael Wu, and members of the Lin lab for discussions and comments on the manuscript. This study was supported by National Institute of General Medical Studies grant R35GM162089 to KZL; R35GM147125 and Brotman Baty Institute Pilot Award to MS.

## Author Contributions

YZ and KZL conceived the idea. YZ implemented the method and performed the analyses. CF and MS processed the Wohlschlegel dataset. All authors discussed the results and wrote the paper together.

