## Supplementary Note for "SCOPE: Localizing fate-decision states and their regulatory drivers in single-cell differentiation"

#### S1 Rationale and discussion of conformal inference

Conformal inference, originally introduced by Vovk et al. [33], provides a framework for uncertainty quantification by determining an empirical quantile threshold from a calibration set to construct prediction intervals for new samples. This principle has been extended across various machine learning tasks [31, 3, 15]. In practice, a widely used implementation is split conformal inference [1, 15], which operationalizes this framework by randomly partitioning the labeled data into a training set and a calibration set. The prediction model is first fit on the training set, and conformal (nonconformity) scores are then computed on the calibration set to estimate the quantile threshold. Compared with full conformal inference, this approach is substantially more computationally efficient, as it requires training the model only once rather than recomputing predictions for each candidate label or test point.

The mathematical validity of conformal prediction relies on the assumption of exchangeability between the calibration data and the test data. Under this assumption, the probability that the true label $Y_{n+1}$ is contained in the prediction set $\mathcal{C}(X_{n+1})$ is guaranteed to be at least $1-\alpha$. In the context of SCOPE, we define non-conformity scores based on the fate probabilities produced by our random forest classifiers, using the iterative recruitment process to provide the necessary pseudo-labels for calibration. As we mentioned in the "Methods" section, although we do violate this exchangeability assumption due to our iterative recruitment of unlabeled cells by pseudotime, we deploy an entropy balancing procedure to mitigate the impact of the distribution shift (see "Entropy balancing to handle distribution shift").

A known property of standard conformal inference is that it provides *marginal* coverage, meaning the $1-\alpha$ guarantee holds on average over the distribution of all cells. This is in contrast to *conditional* coverage, which would guarantee $1-\alpha$ for every specific point in the latent space. In the SCOPE framework, the marginal coverage guarantee is sufficient for our biological goals because biological discovery in single-cell systems fundamentally relies on identifying stable population-level trends rather than cell-specific-level diagnostics. Specifically, in developmental biology, the objective is to characterize the behavior of cell lineages and transition states. Since branchpoints are identified by analyzing signals across thousands of cells, marginal coverage aligns directly with the goal of resolving "consensus" transcriptomic regions of fate-ambiguity. Additionally, with datasets typically containing $10^3$ to $10^5$ cells, a $(1-\alpha) \times 100\%$ marginal guarantee ensures that the vast majority of prediction sets are correctly calibrated. This high volume of valid sets provides more than enough statistical power to infer precise boundaries for fate specification and regulatory lags without requiring the stricter constraints of conditional coverage.

**Specific conformal inference methods that are related.** Two works leveraging conformal inference are noteworthy, in relation to SCOPE. First, conformal inference has recently been used in spatial transcriptomics analyses to predict a gene's expression that is not directly measured in the spatial assay's panel [30]. This application of genomic analyses has a different biological goal than SCOPE, but it demonstrates that the statistical framework of conformal inference is currently being adopted in emerging single-cell analytical workflows. Second, recent work has also explored incorporating unlabeled data into conformal prediction frameworks. For example, CSForest [9] treats unlabeled samples as a pseudo-class and trains a semi-supervised random forest to construct conformal prediction sets under distribution shift. In contrast, SCOPE leverages semi-supervised learning differently by propagating predictions along the cellular manifold and iteratively recruiting cells along pseudotime to study developmental trajectories.

### S2 Qualitative comparison between SCOPE and other methods

#### S2.1 Comparison among methods to localize branchpoints

**Graph abstraction over partitions (cluster-to-cluster topology): PAGA.** PAGA [39] represents differentiation as a coarse-grained graph of partitions (clusters/metacells), where edges reflect statistical connectivity between partitions in the underlying kNN graph. Here, a "branchpoint" is not a particular cell but a partition represented as a node with multiple high-confidence connections. The mathematical principle is graph abstraction with edge-confidence. In terms of uncertainty, PAGA provides edge confidence/connectivity values that act like an internal notion of reliability for the abstracted topology, but it is not typically framed as formal hypothesis testing for a branchpoint.

**Explicit backbone learning as a principal graph / tree (discrete branching nodes in a fitted trajectory): Monocle, STREAM, Slingshot, VITAE.** These methods posit that the data lie near a low-dimensional backbone (often a tree or graph), and they fit that structure directly or fit curves constrained by a tree. Monocle [32, 5] and STREAM [6] learn an explicit principal graph or principal tree, and branchpoints are internal graph nodes where degree exceeds 2 (often interpreted as a local splitting point, mapped back to a set of nearby cells). Alternatively, Slingshot [29] models branchpoints at the cluster level: it builds a minimal spanning tree (MST) over cluster centers, and a branchpoint is an internal MST node where lineages diverge, with simultaneous principal curves sharing structure near the split. VITAE [8] is similar in spirit but probabilistic: it uses a generative latent-state framework with an explicit graph-structured set of states/transitions, so branchpoints correspond to branching vertices in the inferred backbone, and the cells mapping to these states are deemed the branchpoints. Conceptually, all these methods treat branching as the topology of a learned backbone. Uncertainty is mostly handled implicitly (regularization, model selection, stability to parameter choices); these methods usually do not provide formal statistical tests for branchpoints, though they often offer useful diagnostics (fit quality, stability, or likelihood/regularization trade-offs) that can be probed via resampling in practice.

**Local geometry / embedding-first, branch-detection-second (branchpoints as regions where local structure changes): k-Branches, PHATE.** Here, branching is treated as a local geometric phenomenon rather than a node in a fitted global tree. K-Branches [7] defines branching by how well a cell's neighborhood is explained by multiple local rays/half-lines (with model selection for the number of branches), so a "branchpoint" is naturally a set of cells forming a branching region. PHATE [21] is primarily an embedding method, but explicitly proposes identifying branchpoints via increases in local intrinsic dimensionality. In both cases, the mathematical principle is "local model adequacy": branching is detected when the neighborhood geometry ceases to look one-dimensional. Uncertainty is typically based on model selection (e.g., choosing K and sensitivity to neighborhood size and density) rather than hypothesis testing; users often assess robustness by varying neighborhood parameters or subsampling.

**Fate-probability landscapes (branchpoints as decision regimes with low bias / high uncertainty): Palantir, CellRank, CoSpar.** These methods shift the definition from "a topological split" to "a decision regime," in which cells have meaningful probability mass over multiple terminal fates. The mathematical core is a stochastic transition model (typically Markovian on a cell graph): Palantir [28] and CellRank [14] compute absorption/fate probabilities to terminal cell types, and branchpoints are naturally regions of high multilineage potential, often summarized by high entropy or low fate bias. CoSpar [34] similarly builds a transition map across time points (optionally constrained by lineage barcodes) using smoothing and sparsity assumptions; branch/decision regions are where inferred progenitor states remain low-bias among downstream fates and where fate boundaries emerge in the transition structure. These methods generally identify cells along a continuum of how specified to a particular fate, but it is often difficult to define a strict threshold on how far along it a cell needs to be to be labeled as "fate specified." For uncertainty, these are the most naturally probabilistic: they output fate probabilities and derived uncertainty measures, but those are model-based uncertainties rather than formal tests of a branchpoint null hypothesis; robustness is commonly evaluated by sensitivity to kernel construction, terminal-state definition, or bootstrap-style perturbations.

**Lineage-tracing anchored reconstruction (branchpoints as ancestral lineage nodes, possibly unobserved): CARTA.** CARTA [26] is conceptually different because branching is defined primarily in the lineage tree inferred from heritable clonal barcodes, with expression used to annotate/interpret states. A branchpoint corresponds to an ancestral progenitor node in the reconstructed lineage tree, and "branchiness" can be quantified via descendant diversity (potency). The key mathematical principle is reconstructing lineage relationships from barcode similarity/mutation history rather than inferring a manifold backbone from expression alone. The branchpoint object is therefore an (often unobserved) progenitor node mapped to the most plausible transcriptomic states. Uncertainty is largely about the quality of lineage reconstruction (barcode noise, missing intermediates, sampling), and while one can assess the stability of inferred trees, CARTA is typically not presented as performing hypothesis tests for branchpoint existence.

#### S2.2 Comparison among methods to localize epigenetic priming

While various computational workflows have been developed to quantify this phenomenon using single-cell multi-omic data, they diverge in their mathematical framing of priming. Below, we summarize major categories of computational approaches for assessing epigenetic priming and how SCOPE complements each category to build a more holistic understanding of this mechanism.

**Power for predicting the fate.** In this view, priming is defined by predictive asymmetry: early chromatin-derived features (e.g., accessibility patterns or TF-motif activity) can predict later fate outcomes more accurately than RNA-based features alone. This perspective is particularly appealing when the sequencing protocol can directly measure the fate outcome, for example, by lineage tracing [36, 11]. These lineage-tracing workflows use heritable clonal barcodes to demonstrate that early-stage chromatin states are more informative than transcriptomes for predicting the eventual fate of a cell clone. SCOPE is most similar to these workflows, since it also assesses how predictive the ATAC modality is in determining a cell's fate, compared to the RNA modality. However, unlike existing workflows, SCOPE additionally localizes the specific loci (either enhancer regions or genes) that contribute the most towards this prediction. In this sense, SCOPE is spiritually most similar to FateID [10], but as our results demonstrate in Fig. S2, the underlying methodology of SCOPE and FateID is different.

**Forecast of cellular shifts across the entire transcriptome.** In this view, a cell is primed if its chromatin accessibility profile already encodes information about where it is headed in gene-expression space [19, 20]. Methods in this category typically learn a mapping from accessibility to expression, then visualize priming as a vector field or directional tendency in an embedding: chromatin effectively provides a "look-ahead" of transcriptional state. This discrepancy between the two modalities serves as a vector for future transcriptional change. Conceptually, this approach integrates information across broad genomic tiles, capturing the collective influence of multiple enhancers and the overall "readiness" of a gene locus. Unlike SCOPE and other locus-specific models, priming in this framework does not natively pinpoint specific regulatory elements or genes that are primed; rather, it is primarily treated as a global property of the cell state.

**Changes in chromatin accessibility preceding RNA expression at specific loci.** In this view, priming is defined as a temporal offset between chromatin accessibility and gene expression: enhancers regulating a specific locus become more accessible earlier than increases in transcription [41, 12]. Broadly, this is quantified by comparing when accessibility and expression signals "activate" along pseudotime, often through peak–gene associations or other feature summaries. Notably, these methods do not directly assess a cell's fate but rather quantify the temporal relationship between enhancers and genes. Certain computational methods, such as MultiVelo [16], further model ordinary differential equations that explicitly relate the rate of change in chromatin accessibility to the rates of change in the abundance of spliced and unspliced RNA transcripts. Similar to SCOPE, these workflows localize biological signals to specific loci. However, these existing workflows do not directly provide insight into the fate-specification mechanisms without further investigations on which genes are critical for determining a cell's fate. Nonetheless, these methods are synergistic and complementary: SCOPE assesses which loci are critical for fate specification, while existing workflows assess which enhancers forecast changes in transcription. Together, these methods provide a holistic understanding of fate-specification mechanisms.

### S3 Rationale for random forest as opposed to deep-learning classifiers

SCOPE requires a fate classifier that remains stable under an atypical training regime: labels are initially available only for a small set of confidently differentiated (terminal) cells, and the method then iteratively "recruits" earlier cells in pseudotime, using predictions from one iteration as pseudo-labels for subsequent iterations. Consequently, early iterations operate in a pronounced small-sample, class-imbalanced setting, and any instability or overfitting in these early predictions can propagate and amplify across iterations, ultimately distorting the inferred fate-uncertainty landscape that underlies both branchpoint localization and priming analyses.

In principle, modern deep-learning classifiers can be highly expressive and achieve strong performance when ample labeled data are available. However, in our setting, we found them poorly suited to this specific iterative task: the initial labeled set is intentionally small and skewed toward terminal fates, and the model is repeatedly retrained as the labeled set grows. In this regime, we found that deep learning models were often sensitive to initialization, optimization hyperparameters, and class imbalance, leading to unstable probability rankings and, in early rounds, occasionally overly confident predictions. Because SCOPE's iterative recruitment turns these predictions into downstream training targets, such variability is not "washed out" by later iterations; instead, these hallucinated signals can amplify and quickly propagate throughout later recruitment iterations.

We therefore use random forests as the base classifier because they offer a more reliable bias–variance tradeoff in small-sample, imbalanced, and iteratively self-training contexts. This modeling choice is also reflected in other work, such as CellTag-multi [11]. First, random forests are comparatively straightforward to tune with well-established defaults (e.g., number of trees, feature subsampling, depth constraints, and class weighting), and their performance is typically less sensitive to optimizer dynamics than gradient-based deep networks. Second, the bagging principle averages across many trees to substantially reduce prediction variance. This is particularly important in SCOPE because the ordering of fate probabilities determines the composition of the conformal prediction set. In practice, this variance reduction translated to markedly more consistent fate rankings across resamples and across iterations, improving the stability of recruitment and, ultimately, the reproducibility of the inferred branchpoint regions or epigenetic priming signatures.

Finally, we distinguish SCOPE's objective from traditional cell-type classification. While standard single-cell workflows often focus on identifying the immediate identity of transitory cell states, SCOPE is designed to learn each cell's terminal fate potential. In this framework, intermediate progenitors are treated as unlabeled observations whose latent 'true' labels are the mature terminal cell types they are destined to reach. By prioritizing terminal fates over intermediary identities, the model avoids the instability associated with defining discrete boundaries for short-lived cell types and instead provides a more robust interrogation of the continuous landscape of lineage specification.

### S4 Data processing details

**Cell filtering.** We relied on the annotations and count matrix from the original publications, in which the original authors clustered cells and annotated cell populations using marker genes, possibly after applying doublet-detection to remove doublets and ambient-RNA-detection to remove ambient counts. We refer to the original publications for details. Additionally, we removed rare cell populations from our analysis in this paper, since these cell populations have too few cells for any computational method to reliably recover their differentiation patterns.

- **Weinreb dataset**: We removed plasmacytoid dendritic (pDC) and lymphoid cells in our analysis since there are too few cells of these types.
- **Setty dataset**: We didn't remove any cell types.

- **Persad dataset**: We removed CLP and pDC cells because they belong to a lineage distinct from the terminal fates considered for our biological-replicate analysis (Erythroid, Monocyte, DC) [27, 25].
- **Wohlschlegel dataset**: We excluded MuG and BIP cell populations from downstream analyses due to their limited number of cells.

**Selecting highly variable features and denoising count data.** We describe how we preprocessed the RNA modality for each dataset.

- **Weinreb dataset**: We apply a log-normalization (normalizing by sequencing depth of each cell) in accordance with how the original publication analyzed the data. We wanted to apply the same normalization, since the original publication reported detectable clonal relationships in gene expression, and we wanted to use those results as motivation for our in-depth analyses via SCOPE.
- **Setty dataset**: We first used `Scanpy`'s `scanpy.pp.highly_variable_genes()` function to identify 2000 highly variable features [38]. Then, we denoised the gene expression using a customized VAE-based [13] imputation method heavily based on scVI [18]. Specifically, we trained the scVI model with a latent dimensionality of 10. Both the encoder and decoder consisted of two hidden layers with 128 units per layer. The weight on the KL divergence term was set to 0.001. The primary reason we implemented our own VAE is twofold: 1) scVI uses a linear warm-up schedule for the KL divergence penalty, while we used a fixed small value of 0.001, and 2) we explicitly use the observed sequencing depth of the cell as the library size, as opposed to estimating the library size with another neural network. In practice, the performance of scVI and our VAE was very similar, but empirically, we found it easier to tune our VAE.
- **Persad dataset**: The preprocessing for this step was done in the same way as in the Setty dataset.
- **Wohlschlegel dataset**: We also used `Scanpy`'s `scanpy.pp.highly_variable_genes()` function to identify 2500 highly variable features [38] . Then, in accordance with the original publication [37], we log-normalize the data with a pseudocount of 0.1.

We describe the preprocessing steps for the ATAC modality in each multiome dataset.

- **Persad dataset**: We started with the preprocessed counts from the ATAC modality generated by the original authors. Then, we retained peak regions (i.e., features) with nonzero values in at least 15 cells. We then applied PeakVI [2] to impute chromatin accessibility using default model settings. Specifically, the number of hidden units per layer (`n_hidden`) was set to the square root of the number of genomic regions, and the latent dimensionality (`n_latent`) was set to the square root of `n_hidden`. The encoder and decoder neural networks each consisted of two hidden layers (`n_layers_encoder` = 2; `n_layers_decoder` = 2).
- **Wohlschlegel dataset**: We reprocessed the ATAC data from the original publication to filter out doublets and improve peak-calling quality. Specifically, doublets were removed from the ATAC-seq data using Scrublet [40] via the `scrublet.scrub_doublets()` function with default parameters. Then, the remaining cell's fragments were aligned using the hg38 genome as the reference, and peak calling was performed using MACS2, which was found to be more effective than other methods for this dataset. Afterwards, we filtered chromatin accessibility peaks by retaining features with nonzero values in at least 15 cells. The finalized ATAC preprocessing for this dataset follows the default model settings for PeakVI [2], where the number of hidden units per layer was set to the square root of the number of genomic regions, and the latent dimensionality was set to the square root of the number of hidden units. Both the encoder and decoder neural networks consisted of two hidden layers. These processed features were then integrated with pseudotime estimates derived from Palantir analysis of the ATAC-seq data to determine the recruitment order for the SCOPE framework.

**Pseudotime estimation.** We primarily use Palantir [28] to estimate the pseudotime for all the cells. This is needed for SCOPE to bin cells into their recruitment iteration, with cells in earlier recruitment iterations being "closer" to terminal differentiation, and cells in the last recruitment iteration being the most "progenitor-like." To apply Palantir, we need to determine the number of principal components and select specific cells as representatives of the terminal fates.

- **Weinreb dataset**: We first performed PCA using 2000 highly variable genes and constructed a diffusion map from the top 30 principal components. To determine the root cells, we proceeded cell type by cell type. For each cell within a given cell type, we computed its average distance in diffusion map space to undifferentiated cells and selected the cell with the maximum average distance as the root. To define lineage endpoints, we computed Euclidean distances in diffusion eigen-space for the eight lineages found at Day 6, selecting the cell with the maximum average distance from the undifferentiated Day 2 population as the terminal cell types for each lineage. Finally, we ran `palantir.run_palantir()` to obtain pseudotime estimates.

- **Setty dataset**: We followed the pipeline described in the Palantir tutorial notebook (`https://github.com/dpeerlab/Palantir/blob/master/notebooks/Palantir_sample_notebook.ipynb`). Specifically, we normalized the count matrix, embedded the cells in a diffusion map using the top 50 PCs, and used the terminal cell barcodes provided.

- **Persad dataset**: We followed the workflow provided in the SEACells tutorial (`https://github.com/dpeerlab/SEACells/blob/main/notebooks/SEACell_tf_activity.ipynb`). Specifically, we normalized the count matrix, embedded the cells in a diffusion map using the top 30 PCs, and used the terminal cell barcodes provided.

- **Wohlschlegel dataset**: We applied Palantir to the ATAC-seq data to obtain pseudotime ordering. Specifically, after forming the peak-count matrix, we applied Latent Semantic Indexing (LSI) to normalize the counts and computed the top 16 components, dropping the first one due to its correlation with sequencing depth. Then, we embedded the cells into a diffusion map using these components. The terminal cells for each modality's analysis were chosen using the following procedure: The selected terminal cell for each terminal cell type had to be labeled as that cell type via marker genes, and the terminal cell was selected as the cell with the maximum (or minimum) value in the diffusion component that was enriched for that particular cell type. After using Palantir on both the RNA and ATAC modalities, we chose the ATAC pseudotime estimates because they provide a smoother continuum from multipotent cells (MPCs) to all terminal cell types.

**Selecting terminal cell types.**

- **Weinreb dataset**: After removing plasmacytoid dendritic (pDC) and lymphoid cells, there were 8 terminal cell types including: monocytes (Mono), neutrophils (Neu), basophils (Baso), mast cells (Mast), erythroid cells (Ery), megakaryocytes (Meg), eosinophils (EOS), and CCR7-positive dendritic cells (DC).

- **Setty dataset**: To determine the terminal-state clusters, which are required by our method, we performed Louvain clustering with resolution 0.99 and assigned the clusters containing these three annotated cells to their corresponding terminal identities.

- **Persad dataset**: The dataset already included cell type annotations, of which we selected conventional dendritic cell (DC), monocytes (Mono), and erythroid cells (Ery) as the terminal cell types.

- **Wohlschlegel dataset**: The dataset already included cell type annotations, of which we selected Retinal ganglion cells (RGC), horizontal cells (HRZ), and cone cells (CON) as the terminal cell types.

### S5 Pseudocode of SCOPE

See Algorithm 1 to see the pseudocode of SCOPE, which reflects the schematic shown in Figure 1A-D.

**Algorithm 1** SCOPE: Iterative recruitment and conformal prediction

**Input:** Single-cell expression matrix $X \in \mathbb{R}^{n \times p}$ with terminal cell types $\mathcal{Y}_1, \dots, \mathcal{Y}_m \subset \{1, \dots, n\}$; pseudotime estimates $\{t_i\}$.
**Input:** Parameters: miscoverage level $\alpha$ (default: 0.05); number of recruitment iterations $L$ (default: $\lceil \sqrt{n} \rceil$).
**Output:** Calibrated prediction sets $\mathcal{C}(X_i) \subseteq \{1, \dots, m\}$ for all cells $i \in \{1, \dots, n\}$.

**Step 1: Preprocessing and Manifold Construction**
Normalize and denoise $X$; select highly variable features.
Compute low-dimensional embedding $z_i$ and construct a $k$-nearest neighbor graph $G$.

**Step 2: Iterative Recruitment and Conformal Inference**
Partition pseudotime into ordered intervals $\mathcal{I}_1, \dots, \mathcal{I}_L$ from late to early (terminal to progenitor).
Initialize labeled set $\mathcal{L}$ with terminal cells: $\mathcal{C}(X_i) = \{k : i \in \mathcal{Y}_k\}$ for $i \in \mathcal{L}$.
**for** $\ell = 1$ to $L$ **do**
    Define test set $\mathcal{T}_\ell = \{i : t_i \in \mathcal{I}_\ell\}$.
    Partition labeled set $\mathcal{L}$ into training set $\mathcal{L}_\ell^{\text{train}}$ and calibration set $\mathcal{L}_\ell^{\text{cal}}$.
    Train a multi-label binary random forest (warm start) on $\mathcal{L}_\ell^{\text{train}}$.
    Predict fate probabilities $\widehat{\boldsymbol{\pi}}_i$ for $i \in \mathcal{T}_\ell$.
    Smooth probabilities $\widehat{\boldsymbol{\pi}}_i$ over $G$ via label propagation, yielding $\widetilde{\boldsymbol{\pi}}_i$ for $i \in \mathcal{T}_\ell \cup \mathcal{L}_\ell^{\text{cal}}$.
    Compute weights $w_j$ for $j \in \mathcal{L}_\ell^{\text{cal}}$ to minimize distribution shift between $\mathcal{L}_\ell^{\text{cal}}$ and $\mathcal{T}_\ell$.
    Compute conformity scores $s_j$ for $j \in \mathcal{L}_\ell^{\text{cal}}$ based on $\widetilde{\boldsymbol{\pi}}_j$.
    Determine the entrop-weighted empirical $(1-\alpha)$ quantile $\widehat{q}_\ell$ of scores $\{s_j\}_{j \in \mathcal{L}_\ell^{\text{cal}}}$.
    **for** each test cell $i \in \mathcal{T}_\ell$ **do**
        Construct prediction set $\mathcal{C}(X_i) = \{k \in \{1, \dots, m\} : \text{score}(\widetilde{\boldsymbol{\pi}}_i, k) \le \widehat{q}_\ell\}$.
    **end for**
    Update labeled set: $\mathcal{L} \leftarrow \mathcal{L} \cup \mathcal{T}_\ell$.
**end for**

**Algorithm 2** Downstream: Localizing branchpoints

**Input:** Single-cell dataset $X \in \mathbb{R}^{n \times p}$; prediction sets $\mathcal{C}(X_i) \subseteq \{1, \dots, m\}$; two sets of terminal cell types of interest $\mathcal{Y}_a$ and $\mathcal{Y}_b$; local density estimates $\rho_i$ (default: Mellon).
**Input:** Parameters: Number of nearest neighbors $k$ (default: 30); embedding dimension $d$.
**Output:** A set of cells $\mathcal{B}_{a,b} \subseteq \{1, \dots, n\}$ representing the localized branchpoint between $\mathcal{Y}_a$ and $\mathcal{Y}_b$.

**Step 1: Density Estimation**
Using the low-dimensional embedding $z_i$ (e.g., top $d$ principal components), calculate density $\rho_i$ for all cells.

**Step 2: Density clustering**
Select the subset of cells whose prediction sets contain only the queried fates: $\{i : \mathcal{C}(X_i) \in \{\mathcal{Y}_a, \mathcal{Y}_b, \mathcal{Y}_a \cup \mathcal{Y}_b\}$.
Construct a $k$-nearest neighbor graph on $z_i$ for the selected cells.
Prune the graph such that an edge $i \to j$ is retained only if the prediction set of cell $i$ is a subset of the prediction set of cell $j$ (i.e., $\mathcal{C}(X_i) \subseteq \mathcal{C}(X_j)$), enforcing developmental directionality.
Apply density clustering using the pruned graph and density scores $\rho_i$ to identify clusters of fate-ambivalent cells.

**Step 3: Branchpoint Localization**
Define the branchpoint cell set $\mathcal{B}_{a,b}$ as a subset of the selected cells that has the largest cluster.

**Algorithm 3** Downstream: Localizing TFs bearing epigenetic priming

**Input:** Recruitment iterations from SCOPE (ordered from late to early); fate-specific random-forest classifiers from each iteration; gene-enhancer links; list of TF genes of interest.
**Output:** TFs that display epigenetic priming.

**Step 1: Gather importance trajectories across recruitment iterations.**
For each TF (and optionally for each fate-specific classifier), collect a *trajectory* of variable-importance values across the recruitment iterations:
  (i) the TF's gene-expression importance at each iteration, and
  (ii) the TF's enhancer importance at each iteration.
If the TF has multiple linked enhancer regions, sum their importances first so that each TF has a summed enhancer-importance value per iteration.
Linearly rescale each trajectory (gene and summed enhancer) to lie in $[0, 1]$.

**Step 2: Smooth each trajectory with a unimodal fit.**
Independently fit a unimodal regression to:
  (i) the rescaled TF gene-importance trajectory across iterations, and
  (ii) the rescaled TF summed enhancer-importance trajectory across iterations.
For each unimodal fit, record the recruitment iteration at which the fitted curve attains its maximum:
  (i) the apex iteration for the gene-expression trajectory.
  (ii) the apex iteration for the summed enhancer trajectory, and

**Step 3: Declare priming based on which apex happens first.**
Label the TF as epigenetically primed if the summed enhancer apex occurs at least one recruitment iteration before the gene-expression apex, respectively.

### S6 Implementation details of SCOPE

#### S6.1 Details of multi-class random forest classifier

We use the `scikit-learn` implementation of random forests for binary classification via `sklearn.ensemble.RandomForestCla`[illegible] Each classifier is initialized with 100 decision trees and uses class-balanced weighting (`class_weight = "balanced"`) to account for class imbalance. Tree splits are selected by minimizing Gini impurity (`criterion = "gini"`). The minimum number of samples required to split an internal node is set to 2 (`min_samples_split = 2`), and the minimum number of samples required at a leaf node is set to 1 (`min_samples_leaf = 1`). We employ a warm-start strategy during training, retaining previously learned trees while incrementally adding additional trees. By default, 50 new trees are added at each training iteration.

#### S6.2 Details on branchpoint localization and visualization of branchpoints

When localizing the branchpoints, we use the following number of principal components, $d$, for each dataset: Weinreb ($d = 30$), Setty ($d = 35$), Persad ($d = 30$), and Wohlschlegel ($d = 30$). This was based on assessing the scree plot and ensuring that the clustering conformed to the density heatmap when visualized, as in Fig. S3. We also used these latent dimensions when estimating the density via Mellon.

To visualize the branchpoint along a density-pseudotime plot, as shown in Fig. 3f,h in the main text, we perform the following procedure. Suppose we wanted to focus on the branchpoint between erythroid and megakaryocyte fates.

- **Cell Selection**: We included all initially unlabeled cells whose SCOPE prediction sets contained either the erythroid or megakaryocyte terminal fates.
- **Ridge Identification**: To prevent overplotting while preserving the lineage backbone, we filtered for cells residing within the "upper ridge" of the density along this differentiation process. This ridge was determined by regressing cell density against pseudotime (via smoothing spline) after computing

the highest density within sliding windows along pseudotime. We selected cells within a band of this smoothing spline.

- **Aesthetics**: Point sizes correspond to the cardinality of the SCOPE prediction sets, providing a visual measure of "poise" along the trajectory.
- **Branchpoint Annotation**: The horizontal bar indicates the interquartile range (25th–75th quantile) of pseudotime for the localized branchpoint, with associated cells highlighted to emphasize their accumulation within a high-density, fate-ambivalent checkpoint.

##### S6.3 Details on graph construction for label propagation

The construction of the similarity graph in SCOPE follows a multi-step procedure to ensure local structure preservation and global connectivity.

**Forming the adjacency matrix $W$.** Inspired by the Palantir framework [28], we first construct a globally connected graph using an adaptive Gaussian kernel to represent the similarity between cells:

- **Local scale**: For each cell $x_i$, we estimate a local scale $\sigma_i = \frac{1}{k}\sum_{x_j \in \mathcal{N}_k(i)} d(x_i, x_j)$, where $\mathcal{N}_k(i)$ is the set of $k$ nearest neighbors of $x_i$.
- **Mutual kNN**: We compute the directed kNN graph and retain an undirected edge $(i, j)$ only if it is mutual ($i \in \mathcal{N}_k(j)$ and $j \in \mathcal{N}_k(i)$), yielding a symmetric sparse adjacency $A^{\mathrm{mknn}}$.
- **Adaptive kernel weighting**: For each edge in $A^{\mathrm{mknn}}$, we assign a density-aware Gaussian weight:

  $$w_{ij} \;=\; \exp\left(-\frac{d_{ij}^2}{\sigma_i \sigma_j}\right), \qquad w_{ii} = 0$$

  producing the weighted adjacency $W^{\mathrm{mknn}}$.
- **Connectivity via Minimum Spanning Tree (MST)**: To guarantee a globally connected graph, we compute the MST of the kNN distance graph. Let $W^{\mathrm{mst}}$ denote this MST with edge weights set to half the minimum non-zero weight in $W^{\mathrm{mknn}}$. This ensures connectivity while allowing local mutual-$k$NN edges to dominate.
- **Final graph**: The final sparse adjacency $W$ is the element-wise maximum of the mutual-$k$NN kernel graph and the reweighted MST: $W = \max\{W^{\mathrm{mknn}}, W^{\mathrm{mst}}\}$.

**Normalized adjacency matrix.** Following the construction of $W$, we perform symmetric normalization to stabilize downstream spectral properties [22]:

$$S = D^{-\frac{1}{2}} W D^{-\frac{1}{2}}$$

where $D$ is a diagonal degree matrix with $D_{ii} = \sum_{j=1}^{n} W_{ij}$. This normalization ensures that the resulting graph Laplacian is robust to outliers and variations in sampling density.

(Note: We deliberately do not use this graph constructed for label propagation when we perform density clustering for localizing branchpoints later, since disconnected components of the kNN graph constructed in density clustering are informative when clustering cells. In contrast, label propagation discussed here suffers in performance if the graph has disconnected components since the label information cannot effectively propagate in that scenario.)

#### S6.4 Details on correlating prediction entropy and pseudotime

To numerically validate that the identified branchpoints represent regions of rapid lineage specification, we analyzed the relationship between cellular uncertainty and developmental progression. For a given branchpoint between terminal fates $A$ and $B$, we first quantified the fate uncertainty for each cell $i$ using the Shannon entropy of its predicted fate probabilities $\widetilde{\pi}_i$:

$$H(i) = -\sum_{j=1}^{m} \widetilde{\pi}_{ij} \log(\widetilde{\pi}_{ij}),$$

where $m$ is the number of terminal cell types. We then modeled the relationship between entropy and pseudotime $t_i$ using linear regression:

$$\widehat{H}(i) = \beta_0 + \beta_1 t_i + \epsilon_i.$$

In this formulation, the slope $\beta_1$ represents the rate of change in fate uncertainty as cells progress toward differentiation. A strongly negative slope indicates a rapid resolution of fate-ambivalence into committed singleton sets.

To assess the statistical significance of the observed slope ($\widehat{\beta}_{1,\text{obs}}$) relative to the broader developmental context, we implemented a permutation-based null model. We defined a lineage-specific pool consisting of all cells whose prediction sets were $\{A\}$, $\{B\}$, or the fate-ambivalent set $\{A, B\}$. We then performed 500 iterations of the following procedure:

1. Randomly subsample a set of cells from the lineage pool, equal in size to the number of cells in the estimated localized branchpoint.

2. Fit the linear regression model to the subsampled set and record the resulting slope $\widehat{\beta}_{1,\text{null}}$.

The empirical $p$-value was calculated as the proportion of null iterations where $\widehat{\beta}_{1,\text{null}}$ was more negative than $\widehat{\beta}_{1,\text{obs}}$. This p-value and null distribution were shown in Fig. 3e in the main text.

#### S6.5 Details on differential expression within a branchpoint via TradeSeq

We use `nknots=6` in the `tradeSeq::fitGAM()` function to assess which genes are differentially expressed based on fate bias. Since our goal was to demonstrate biological replicability between the Setty and Persad datasets, we assessed differential expression across all genes in both datasets (rather than focusing only on highly variable genes) to increase the number of genes shared between the two. This method also relies on pseudotime and fate bias, which Palantir already outputs when we learned binned cells into recruitment iterations. We used adjusted p-value cutoffs of 0.05 for both datasets. There are 267 cells at the erythroid–monocyte branchpoint in the Setty dataset and 488 cells at the same branchpoint in the Persad dataset.

#### S6.6 Details on gene ontology analysis

We used the `enrichGO()` function from the `clusterProfiler` [42] package for gene set enrichment analyses and specified `org.Hs.eg.db` as the database.

#### S6.7 Details on log-fold change between fates in a branchpoint

To assess the consistency of molecular drivers across independent datasets, we calculated the gene-level expression bias between the erythroid and monocyte lineages within the identified branchpoints. For each dataset (Setty and Persad), we first identified the set of cells $\mathcal{B}_{Ery,Mono}$ constituting the monocyte-erythroid branchpoint.

These cells were partitioned into two subpopulations based on their estimated fate bias derived from Palantir:

- Erythroid-biased subset: $\mathcal{S}_{Ery} = \{i \in \mathcal{B}_{Ery,Mono} : \pi_{i,Ery} > \pi_{i,Mono}\}$

- Monocyte-biased subset: $\mathcal{S}_{Mono} = \{i \in \mathcal{B}_{Ery,Mono} : \pi_{i,Mono} > \pi_{i,Ery}\}$

For each gene $g$ in the set of highly variable genes, we computed the mean normalized and imputed expression level within each subset:

$$\mu_g^{(Ery)} = \frac{1}{|\mathcal{S}_{Ery}|} \sum_{i \in \mathcal{S}_{Ery}} X_{ig}, \quad \mu_g^{(Mono)} = \frac{1}{|\mathcal{S}_{Mono}|} \sum_{i \in \mathcal{S}_{Mono}} X_{ig}$$

The lineage-specific expression bias, or log-fold change (FC), was then defined as the difference in mean expression:

$$FC_g = \mu_g^{(Ery)} - \mu_g^{(Mono)}$$

Since the input expression matrix $X$ was already log-normalized, this difference corresponds to the log-fold change. We performed this calculation independently for both the Setty and Persad datasets and calculated the Pearson correlation of the resulting $FC_g$ values across all common genes to evaluate the degree of biological replicability (shown in Fig. 4e in the main text). This approach ensures that we are comparing the molecular "signature" of the fate specification itself, regardless of the specific cell counts or global manifold density in either dataset.

#### S6.8 Details on linking enhancers to genes via SEACells

To identify regulatory relationships between chromatin accessibility peaks and target gene expression, we employed the SEACells framework [23]. This approach aggregates single-cell data into "metacells," groups of highly similar cells that represent distinct cellular states, to overcome the inherent sparsity of single-cell multi-omic measurements while preserving granular heterogeneity. By calculating the correlation between peak accessibility and gene expression across these metacells, we defined putative enhancer regions for transcription factors (TFs) analyzed in our epigenetic priming workflows.

**Persad dataset.** For the Persad dataset, we followed the established SEACells multi-omic workflow to define the regulatory landscape of myeloid differentiation via the `SEACells.core.SEACells()` function. We constructed 90 metacells, and the diffusion kernel for metacell construction was computed using the PCA embedding of the RNA modality, with the number of waypoint eigenvectors (`n_waypoint_eigs`) set to 10. To ensure maximum stringency for our deep-dive into erythroid drivers, we defined the enhancer regions for the GATA2 locus (Fig. 5d) by evaluating peak-gene links using a correlation cutoff of 0.3, followed by a Benjamini-Hochberg multiple-testing corrected p-value cutoff of 0.05.

**Wohlschlegel dataset.** In the Wohlschlegel dataset, we applied the SEACells framework to characterize the enhancer-gene associations driving retinal cell-fate acquisition. Here, we constructed 150 metacells, and the diffusion kernel for metacell construction was computed from the PCA embedding of the RNA modality, with 14 waypoint eigenvectors. Following metacell aggregation and peak-gene linking, we defined the cis-regulatory landscape for candidate TFs using a correlation threshold of 0.1, followed by a Benjamini-Hochberg-corrected p-value cutoff of 0.05. This threshold was selected to facilitate a systematic, genome-wide screen of 18 highly variable TFs in the cone lineage, ensuring that the informational apex analysis could capture regulatory poise even among enhancers with subtle accessibility leads.

### S7 Details on simulated data generation

To evaluate the framework's performance in complex developmental scenarios, we generated synthetic single-cell data with a hierarchical branching topology. The simulation consists of three primary stages: latent skeleton construction, high-dimensional manifold projection, and the derivation of a global pseudotime for iterative recruitment.

**Latent skeleton and noise model.** The simulation is grounded in a two-dimensional latent space where cells are sampled along four linear segments to form a tree structure with three terminal fates. The segments connect an initial progenitor state to three endpoints: Fate 1, Fate 2, and Fate 3. A primary bifurcation occurs at the origin, followed by a secondary bifurcation along the path leading to Fates 2 and 3. Individual cell coordinates are sampled uniformly along these segments, with Gaussian noise $\sigma$ applied to coordinates to simulate biological and technical stochasticity.

**High-dimensional feature generation.** To mimic the complexity of single-cell measurement spaces, the two-dimensional latent coordinates are expanded into a 240-dimensional feature space via a linear transformation, with noise added afterwards. This transformation ensures that the underlying branching structure is embedded within a high-dimensional manifold, requiring the framework to recover the developmental topology from global geometric relationships rather than simple 2D proximity.

**Pseudotime and recruitment order.** A global pseudotime $t_i \in [0, 1]$ is assigned to each cell $i$ based on its progress along the hierarchy. The root progenitor is defined at $t_i = 0$, with the first bifurcation resolving as the trajectory progresses. The secondary branch resolves into Fates 2 and 3 at $t_i = 0.5$, with terminal cell types reaching $t_i = 1$. This time-ordering serves as the recruitment sequence for the iterative conformal prediction process, facilitating the propagation of fate labels from mature terminal states back to early progenitors.

### S8 Additional results

#### S8.1 Additional results on simulated data

Our simulated data in Fig. 2 in the main text were not generated by a synthetic single-cell simulator (although this simulation strategy was used in [17]). To ensure that SCOPE works well on existing single-cell simulating workflows, we deployed SCOPE on dyngen [4] following the simulation strategy in [35] (Fig. S1; the blue 'sA' cells differentiate into red 'sC' or purple 'sD' cells). SCOPE estimated the prediction sets with high power, but this figure demonstrates why we did not showcase dyngen's results in the main text – many existing single-cell simulation workflows produce single-cell datasets in which the "signal" is too large and the differentiation pattern is too simplistic to yield a meaningful benchmark.

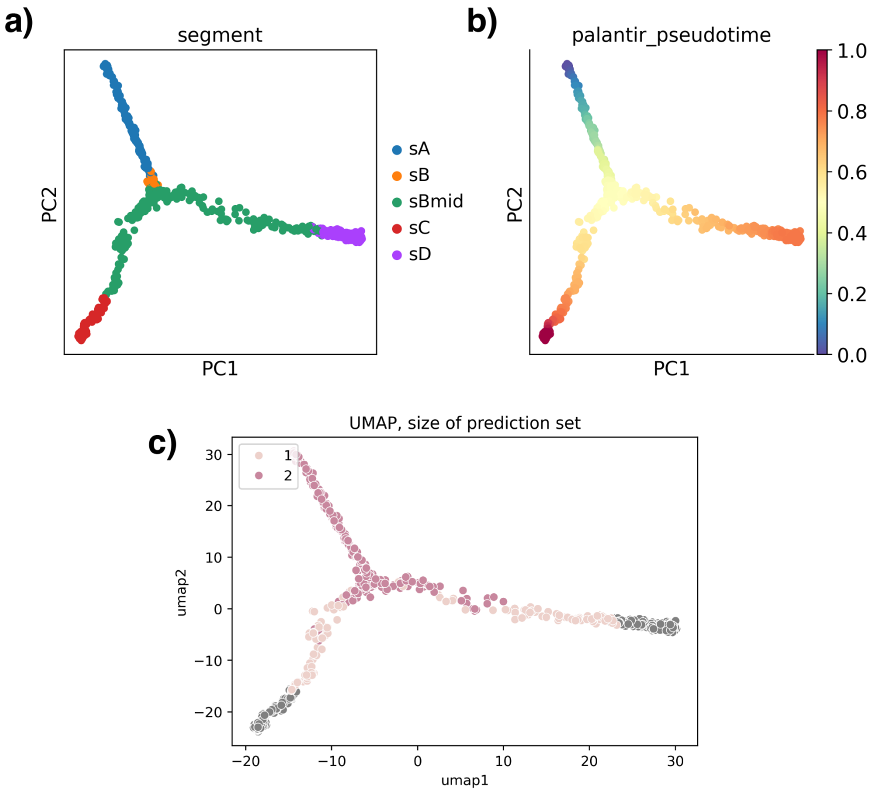


Figure S1: **Result for dyngen simulation dataset. a)** PCA plot showing dyngen simulation dataset. sC and sD are terminal cell types. **b)** PCA plot showing the Palantir pseudotime estimate for the simulation dataset. **c)** UMAP plot showing the conformal result (size of prediction set) of the simulation dataset.

As mentioned in Sec. S2.1, SCOPE is conceptually most similar to FateID [10], but FateID's modeling choices make it quite sensitive to the scRNA-seq data. When we applied FateID to the simulated data shown in Fig. 2a, we observed that FateID's inferred recruitment order deviated substantially from the ground truth (Fig. S2a,b). Moreover, FateID identified only the branchpoint at the intersection between the segments of Cluster 1 and Cluster 7, while failing to detect the branchpoint between the segments of Cluster 1 and Cluster 8 (Fig. S2d-f). This biased branchpoint between Clusters 1 & 7 with Cluster 8 stems from the fact that it is difficult to manipulate FateID's recruitment order, since FateID uses correlation of gene expression to determine a suitable recruiting order. In contrast, since SCOPE takes as input cell pseudotime, it can leverage the most modern pseudotime method to yield more stable performance (where we currently use Palantir [28] throughout the paper). There are also statistical advances in SCOPE over FateID (e.g., SCOPE's use of conformal inference, which provides interpretable, statistically rigorous prediction sets).

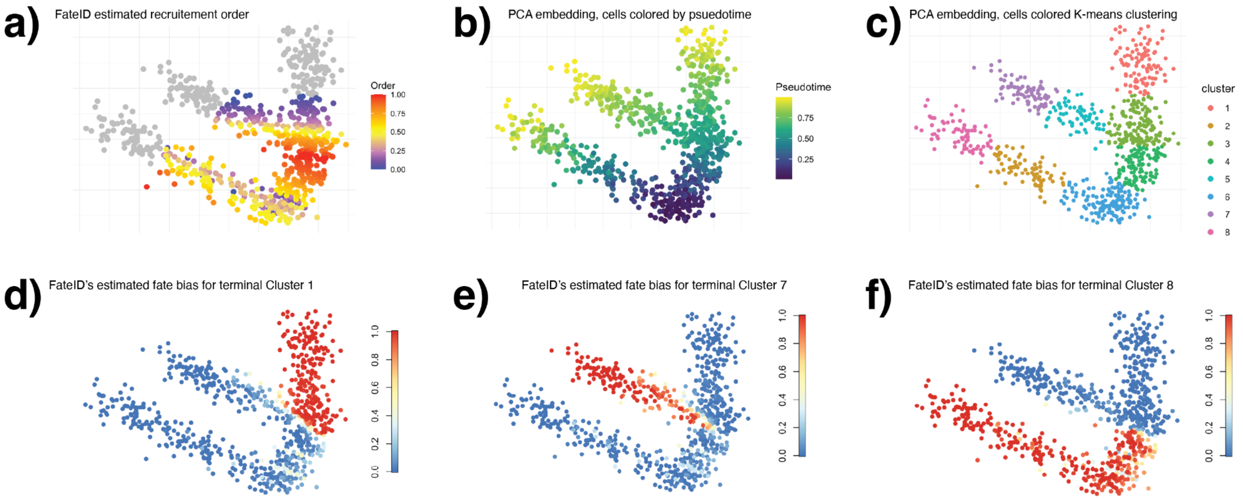


Figure S2: **FateID result for our simulated dataset.** **a)** FateID's recruitment order, for the simulated data shown in Fig. 2a in the main text. **b)** Ground truth recruitment order. **c)** Clustering result for simulation data. **d)** Fate bias for a terminal cell type (Cluster 1). **e)** Fate bias for a terminal cell type (Cluster 7). **f)** Fate bias for a terminal cell type (Cluster 8). **g)** FateID inferred trajectory.

#### S8.2 Additional results on Weinreb dataset

We display additional results for the Weinreb dataset [36]. In Fig. S3, we investigate the impact of applying a density clustering among all the unlabeled cells labeled with {Mono}, {Neu}, or {Mono, Neu} (light blue, light orange, and green, respectively; Fig. S3a). If we labeled all the cells with a prediction set of {Mono, Neu} as the branchpoint, we would yield potentially unsatisfactory results because: (1) there are many different "clusters" of cells with {Mono, Neu} (Fig. S3a), and (2) most of the density of cells with a prediction set of {Mono, Neu} is actually split between two different high density regions (Fig. S3b). Hence, empirically, we found that performing a density clustering is critical. We show the five different density clusters we estimated among the unlabeled cells labeled with {Mono}, {Neu}, or {Mono, Neu} in Fig. S3c-g.

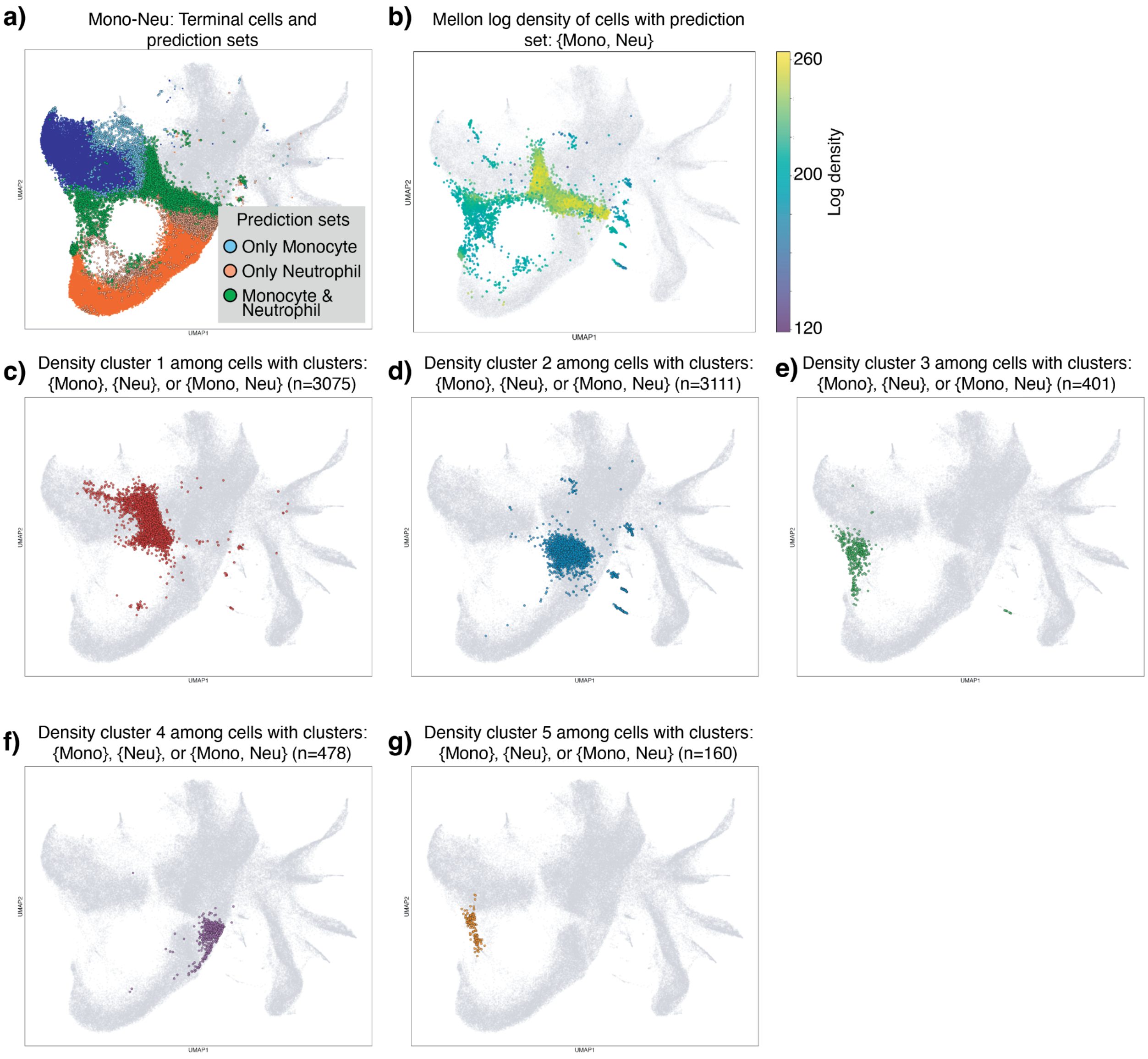


Figure S3: **Density clustering for Mono-Neu branchpoint.** **a)** UMAP of the Weinreb datasets, coloring the terminal monocyte and neutrophils similarly as in Fig. 2e, as well as initially unlabeled cells that were estimated to have a prediction sets of {Mono} (light blue), {Neu} (light orange), or {Mono, Neu} (green). **b)** Estimated density among initially unlabeled cells that were estimated to have a prediction set of {Mono}, {Neu}, or {Mono, Neu}. **c-g)** Each of the five estimated density clusters, among initially unlabeled cells that were estimated to have a prediction set of {Mono}, {Neu}, or {Mono, Neu}.

Since the Weinreb dataset included clonal lineage barcodes, we also used them to assess the "validity" of our estimated Mono-Neu branchpoint. To recap, these clonal lineage barcodes were delivered via lentivirus to each cell prior to the start of the experiment, and were inherited by the original cell's progeny as it divided. This way, we can assess temporal relationships despite single-cell sequencing being destructive. We would expect the "true" branchpoint between monocytes and neutrophils to be highly enriched with clonal lineages where (1) there are a good portion of undifferentiated cells, and (2) a good portion of differentiated monocytes *and* differentiated neutrophils. This is because a true branchpoint should depict cells that are making a fate specification to one fate or another. We first visualized the clones (i.e., clonal lineages) with the largest number of undifferentiated cells as a reference (Fig. S4a). Next, we visualized the 10 clones with the largest number of undifferentiated cells, among all the clones detected within our estimated branchpoint

(Fig. S4b). To clarify, not all the cells in Fig. S4b were located inside the estimated branchpoint. We depicted the number of cells that were within the estimated branchpoint in parentheses on the bottom of the x-axis. We saw that almost every one of the top 10 clones contained cells that differentiated into monocytes (blue) and neutrophils (orange), suggesting that our branchpoint indeed captured cells making a specification to one of these fates. As a reference, we also plotted the 10 clones with the largest number of undifferentiated cells, whereby the only fate in that clonal lineage was either monocytes (Fig. S4c) or neutrophils (Fig. S4d). Comparing Fig. S4c,d to Fig. S4b, we saw that SCOPE's ability to localize the branchpoint was impressive – there were many "pure" (i.e., unipotent) clones much larger than those represented in the estimated branchpoint, and SCOPE did not erroneously select many of these clones.

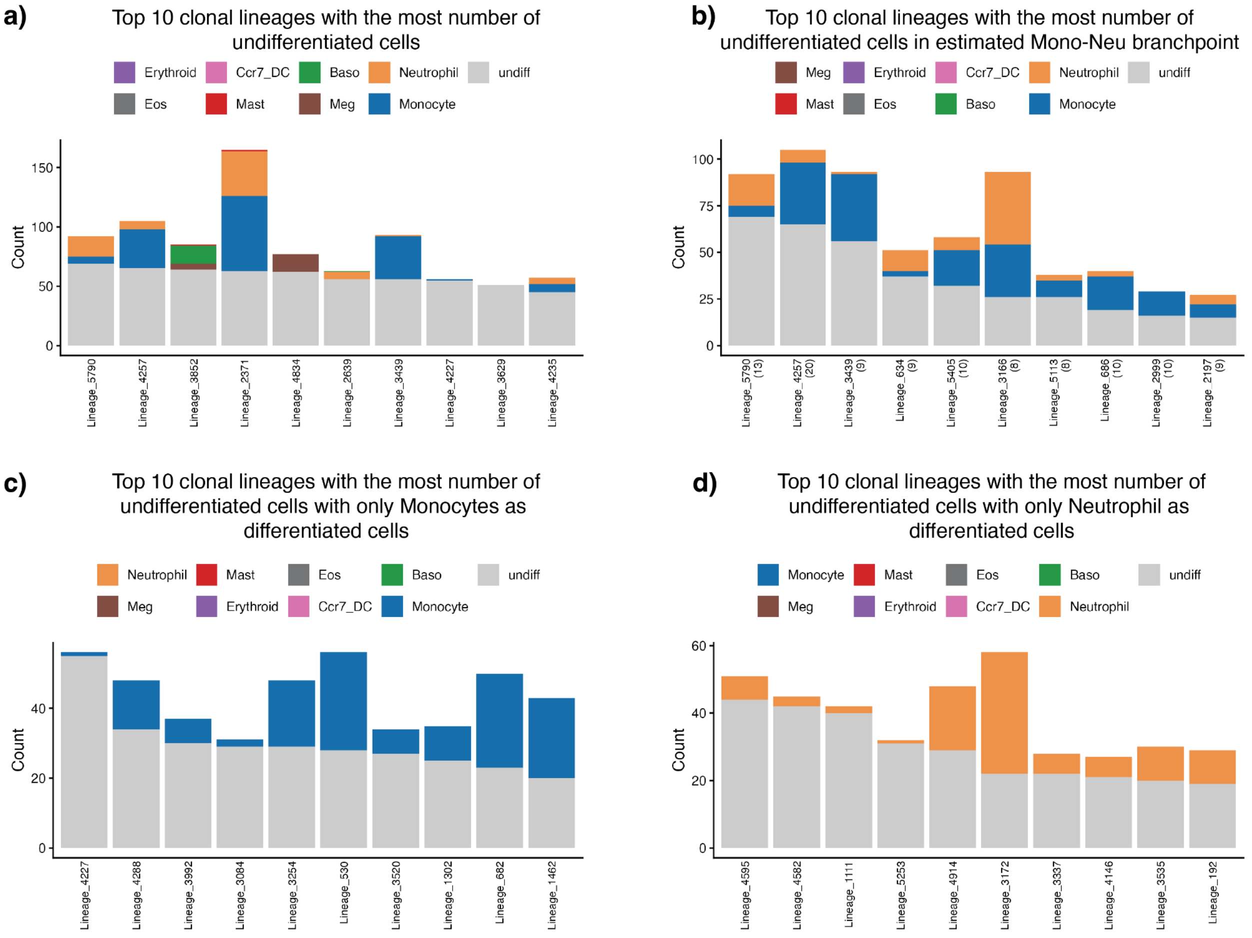


Figure S4: **Cell type composition among clonal lineages in the Weinreb dataset. a)** Composition of the largest 10 clones (i.e., clonal lineages), ranked by the number of undifferentiated cells, across the entire Weinreb dataset. **b)** Composition of the largest 10 clones, ranked by the number of undifferentiated cells, among the clones represented by SCOPE's estimated Mono-Neu branchpoint. The number of parentheses represents the number of cells (among all the cells with a particular clonal barcode) detected to be inside the branchpoint. **c)** Composition of the largest 10 clones, ranked by the number of undifferentiated cells, among all unipotent clones where the only differentiated cell type was monocytes. **d)** Composition of the largest 10 clones, ranked by the number of undifferentiated cells, among all unipotent clones where the only differentiated cell type was neutrophils.

We also investigated other branchpoint between two terminal cell types. In Fig. S5b,c, we plotted the branchpoint between erythroids and basophils and the branchpoint between erythroids and mast cells, respectively. Additionally, we applied the entropy-pseudotime diagnostic on the Mono-Neu branchpoint, analogous to Fig. 3e in the main text (Fig. S5d). The entropy versus pseudotimes for both the Meg-Ery branchpoint analyzed in the main text and the Mono-Neu branchpoint are shown in Fig. S5e,f.

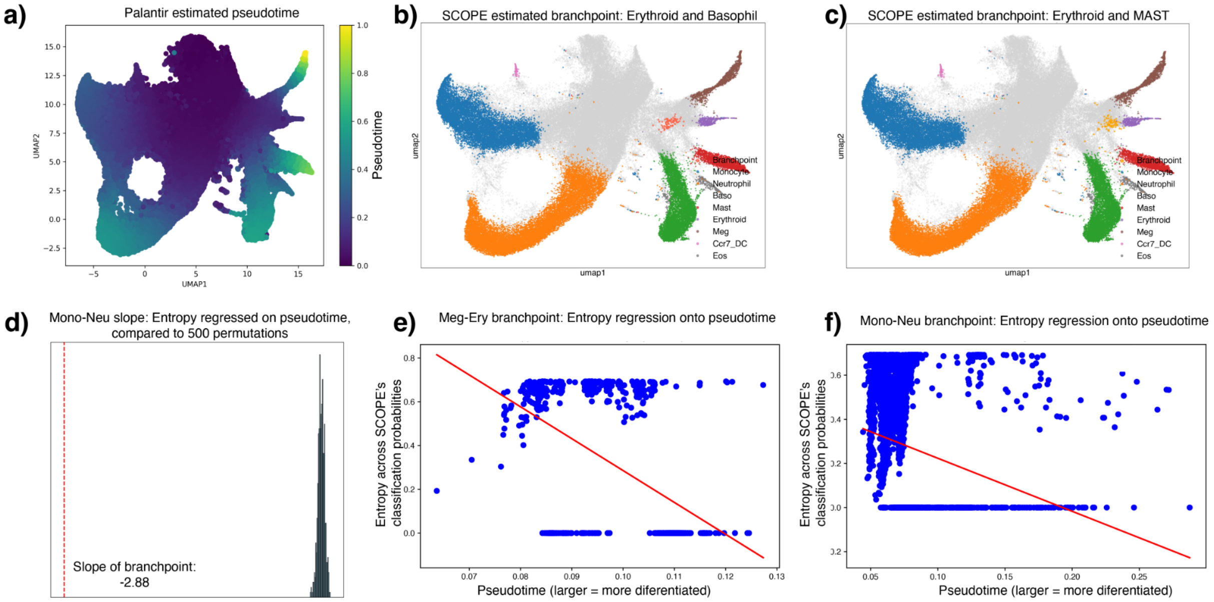


Figure S5: **Additional SCOPE branchpoint and diagnostics for the Weinreb dataset a)** UMAP plot showing the Palantir pseudotime estimate for the Weinreb dataset. **b)** UMAP plot showing Ery-Baso branchpoint. **c)** UMAP plot showing Ery-Mast branchpoint. **d)** Null distribution of slopes for the branchpoint of Mono and Neu, when compared to the estimated slope. **e)** Linear regression between entropy and pseudotime for cells within the estimated Ery-Meg branchpoint. **f)** Linear regression between entropy and pseudotime for cells within the estimated Mono-Neu branchpoint.

While most of the main text focused on branchpoints between two terminal cell types, SCOPE actually allows for much broader investigations. We plotted the number of unlabeled cells for each combination of terminal cell types in their prediction set, across the 18 most-represented prediction sets (Fig. S6). To clarify, 1) the order of cell types in the prediction set does not matter, 2) there were a total of $2^8 - 1 = 255$ unique prediction sets, representing any combination of the 8 terminal cell types, and 3) our number of cells shown in Fig. S6 did not include the originally labeled (terminal) cells. Understandably, the most common prediction set included all 8 terminal cells, representing the most "progenitor-like" cells that have made no specification to any fate. The next most common prediction sets are unipotent or bipotent, reflecting the enriched representation in this dataset for cells that can differentiate into monocytes, neutrophils, or basophils. Interestingly, afterwards, there is a high number of cells that had all terminal cells in their predictions, minus DC cells ($n = 4833$), minus DC cells and eosinophils ($n = 2030$), minus DC cells, eosinophils, and mast ($n = 1570$), and so on. This suggests that a future extension of SCOPE could leverage the abundance of each terminal-cell combination in their prediction sets to reconstruct a potential lineage tree, in a similar spirit to Carta [26]. As we mentioned in Sec. S2.1, this perspective recasts differentiation less as a "specification" to a particular fate, but more as successive "elimination" of the potency to differentiate into particular fates.

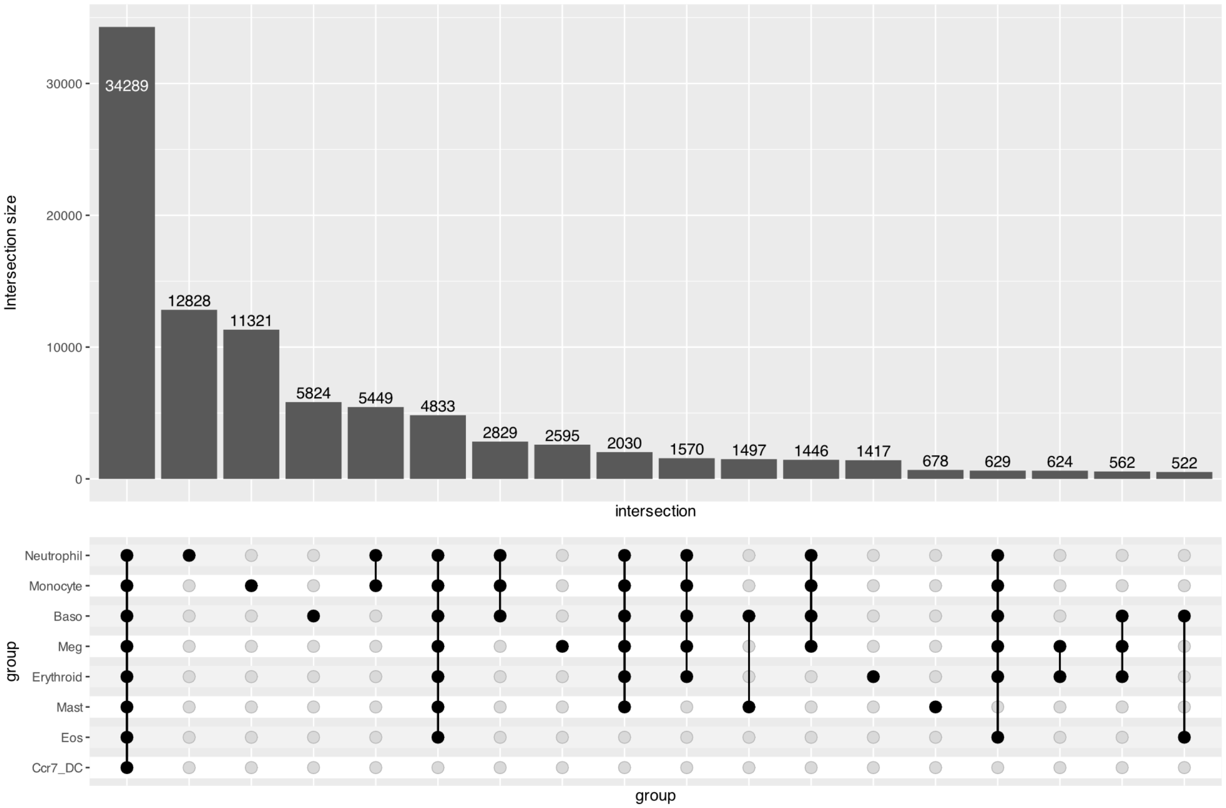


Figure S6: **Upset plot of the top 18 prediction sets among unlabeled cells in the Weinreb dataset.** The horizontal bars depict the number of initially unlabeled cells with a particular combination of terminal cells in their prediction sets, while the dots below depict the particular combination of terminal cells.

We demonstrated that SCOPE's branchpoint localization is not restricted to branchpoints between two terminal branchpoints by demonstrating the estimated branchpoint between {Mono, Neu} and {Baso} (Fig. S7a). This was inspired by recent findings in Carta [26] that suggest that during differentiation, cells either specify their fate to become a basophil or remain bipotent for differentiating into monocytes or neutrophils. We plotted cells with various prediction sets in the rest of Fig. S7, including the three prediction sets involved in estimating this "inner" branchpoint (Fig. S7b) and some of the largest multipotent prediction sets estimated by SCOPE (Fig. S7c-f).

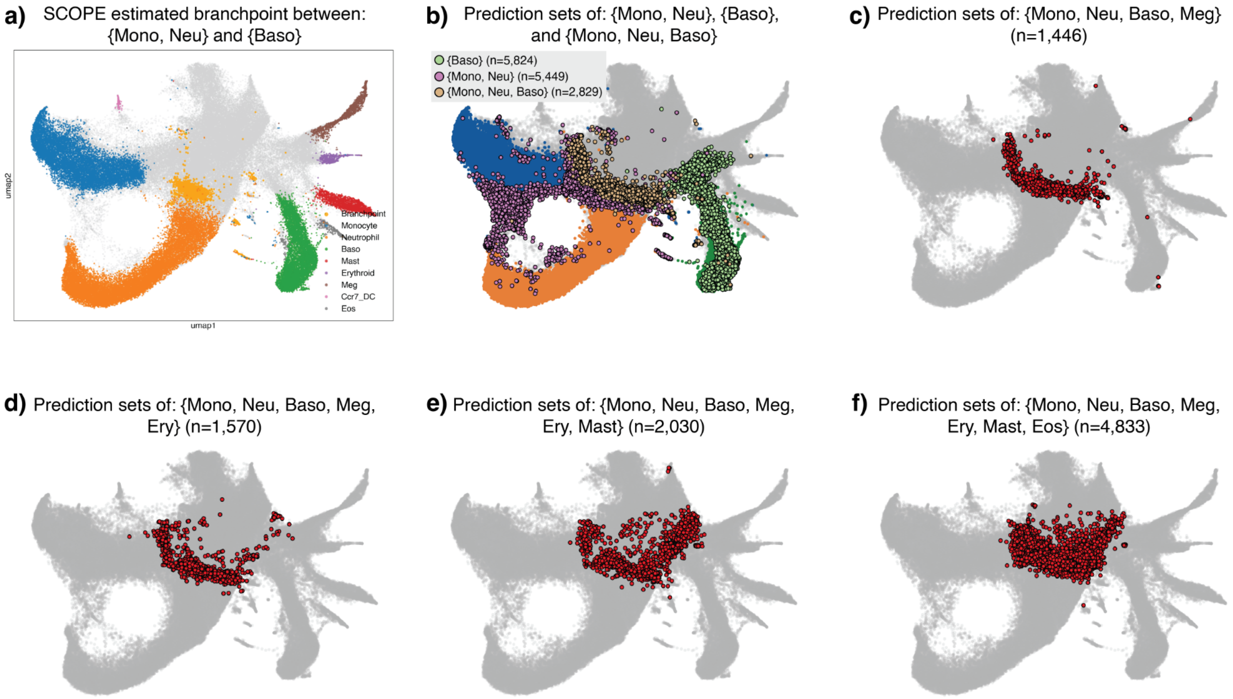


Figure S7: **Multipotent prediction sets in the Weinreb dataset. a)** The "inner" branchpoint between the fates of {Mono, Neu} and {Baso} marked as bright orange. **b)** Cells marked by either their prediction sets being {Mono, Neu} (light pink), {Baso} (light green), or {Mono, Neu, Baso} (light brown). **c-f)** Cells marked by certain multipotent prediction sets.

We next benchmarked other existing methods on the Weinreb dataset. In Fig. S8, we plotted the fate bias estimated by Palantir. While these fate biases looked promising, they quickly diminished to 0 once the gene expression differed from that of the terminal (labeled) cell type. This is primarily because Palantir's fate bias was designed mainly for "bias to a unipotent fate," whereas SCOPE can quantify varying degrees of uni/multipotency via the conformal prediction sets.

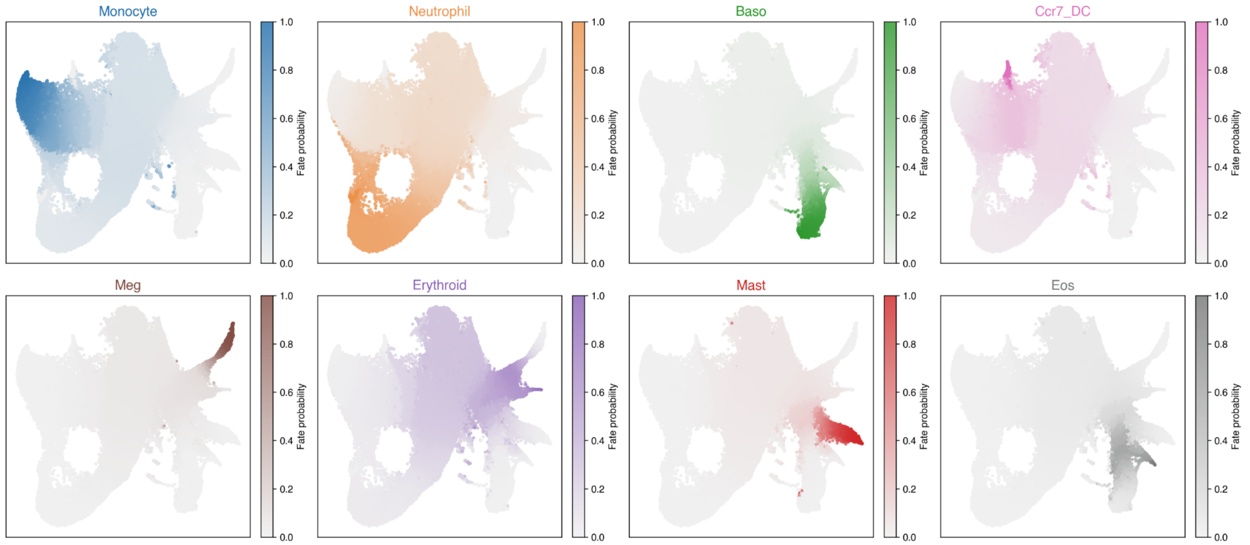


Figure S8: **Fate bias derived by Palantir for 8 terminal cell types in the Weinreb dataset.**

Next, we applied CoSpar [34] to the Weinreb dataset. Unlike Palantir, CoSpar leverages the clonal lineage barcodes, so one would expect a markedly increased localization of branchpoints by leveraging additional information that neither SCOPE nor Palantir has access to. In Fig. S9, we show the fate map estimated by CoSpar. These look similar to Palantir's fate bias (Fig. S8), but arguably are a bit more skewed towards large terminal cell types (mainly monocytes or neutrophils). Overall, these results by themselves were promising, but the main limitation occurred when we tried to localize the branchpoint using CoSpar (Fig. S10). The two limitations we faced were: 1) CoSpar was only able to define fate bias between two terminal cell types, and 2) while certain cells indeed had a "neutral" fate bias (i.e., fate bias of 0.5 with respect to the 2 terminal cell types), there was almost no localization of this neutral fate bias. A high proportion of initially unlabeled cells was deemed fate-neutral, which is unlikely to reflect the underlying biology, as CoSpar's fate bias calculations do not account for the other terminal cell types.

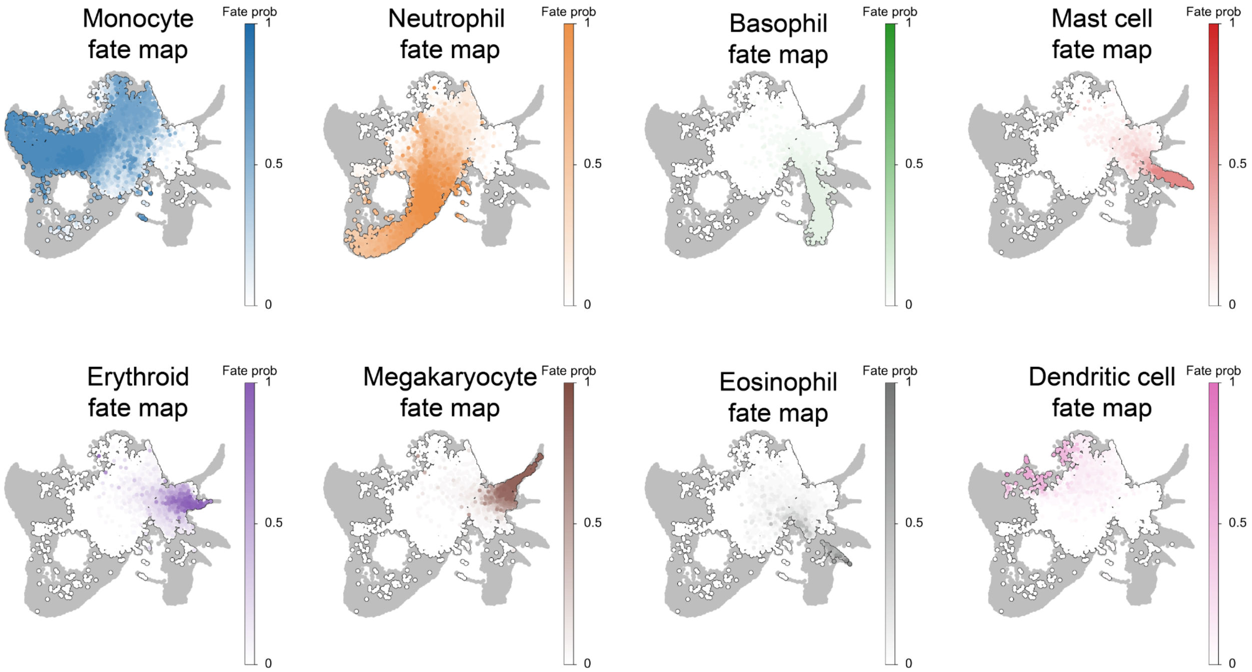


Figure S9: **Fate map estimated by CoSpar on the Weinreb dataset.** CoSpar leverages clonal lineage barcodes when estimating the fate map, information that neither SCOPE nor Palantir uses.

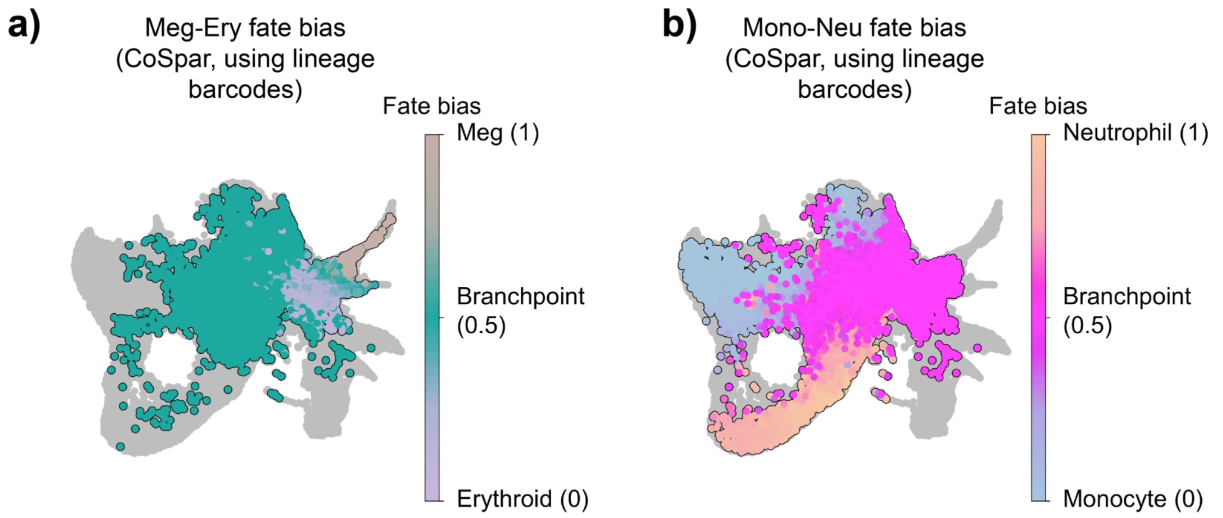


Figure S10: **Fate bias estimated by CoSpar on the Weinreb dataset.** **a)** Fate bias between megakaryocytes (brown) and erythroids (purple), where the "neutral" fate bias cells are colored in green. **b)** Fate bias between neutrophils (orange) and monocytes (blue), where the "neutral" fate bias cells are colored in magenta. Only the initially unlabeled cells are colored (i.e., no terminal cells are colored).

Next, we applied CellRank [14] to the Weinreb dataset, which leveraged the spliced/unspliced counts in an RNA velocity framework. (We credit Pyro-velocity [24] for providing the spliced/unspliced counts.) We saw the RNA velocity flow was quite promising, minus potential "backward" arrows for the differentiation of megakaryocytes (Fig. S11a). However, when we used the estimated transition matrices from CellRank to localize the branchpoint, we were unable to successfully identify any branchpoint within reason. We assessed that this limitation stemmed from a sudden increase in the entropy of the predicted probabilities estimated from CellRank, where many cells had too high an entropy (i.e., the predicted probability of committing to a particular fate was equally distributed across all 8 fates) (Fig. S11b). These cells were high-entropy and biased towards the monocyte and neutrophil fates. In contrast, the entropy of predicted probabilities from SCOPE based on its multi-label random forests increased much more gradually when going from the labeled terminal cells to progenitor-like cells (Fig. S11c). This empirical distinction enabled SCOPE to successfully localize branchpoints.

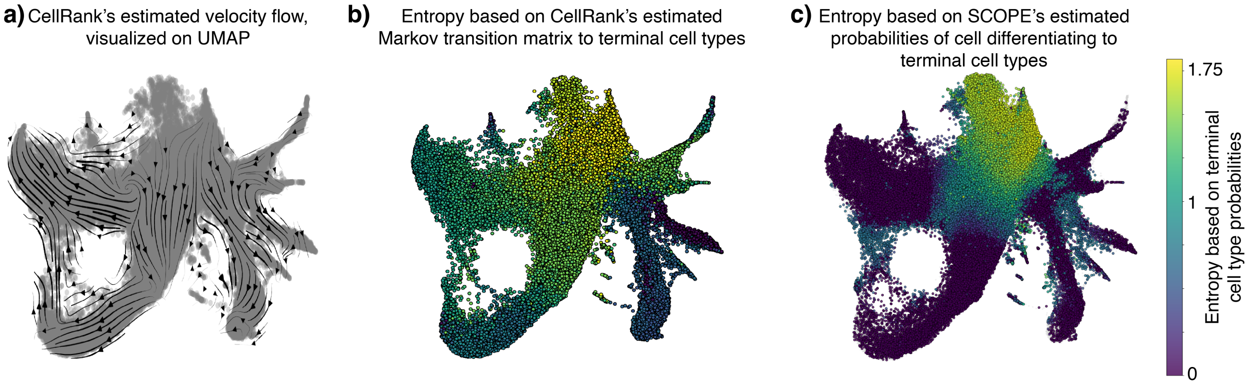


Figure S11: **CellRank results on the Weinreb datasets. a)** RNA velocity flow, as estimated via CellRank by leveraging the relationship between spliced and unspliced counts for each gene. This information is neither used by SCOPE nor by Palantir. **b)** The entropy is based on the predicted probability that a cell differentiates into any of the 8 terminal cells. This calculation was done by leveraging Markov matrix properties, based on the transition matrix estimated by CellRank. **c)** The entropy is based on the predicted probability that a cell differentiates into any of the 8 terminal cells, based on SCOPE's multi-label random forest classifiers.

#### S8.3 Additional results on Setty and Persad datasets

As mentioned in Sec. S4, the Setty dataset lacked cell-type labels. Rather, specific cells were labeled for each of the three terminal cell types (DC, Monocyte, Erythroid). Since SCOPE requires a set of cells representing each terminal cell type, we first performed a clustering on the Setty dataset's gene expression (Fig. S12a). We picked a resolution such that each labeled terminal cell was in its own cluster, and the resulting clusters containing each of the three terminal cells were approximately equal in size. Then, we labeled the terminal cell types as the cluster that contains the labeled terminal cell (Fig. S12b). This yielded the estimated pseudotime (Fig. S12c). Additionally, when localizing branchpoints, SCOPE required the cell densities (Fig. S12d).

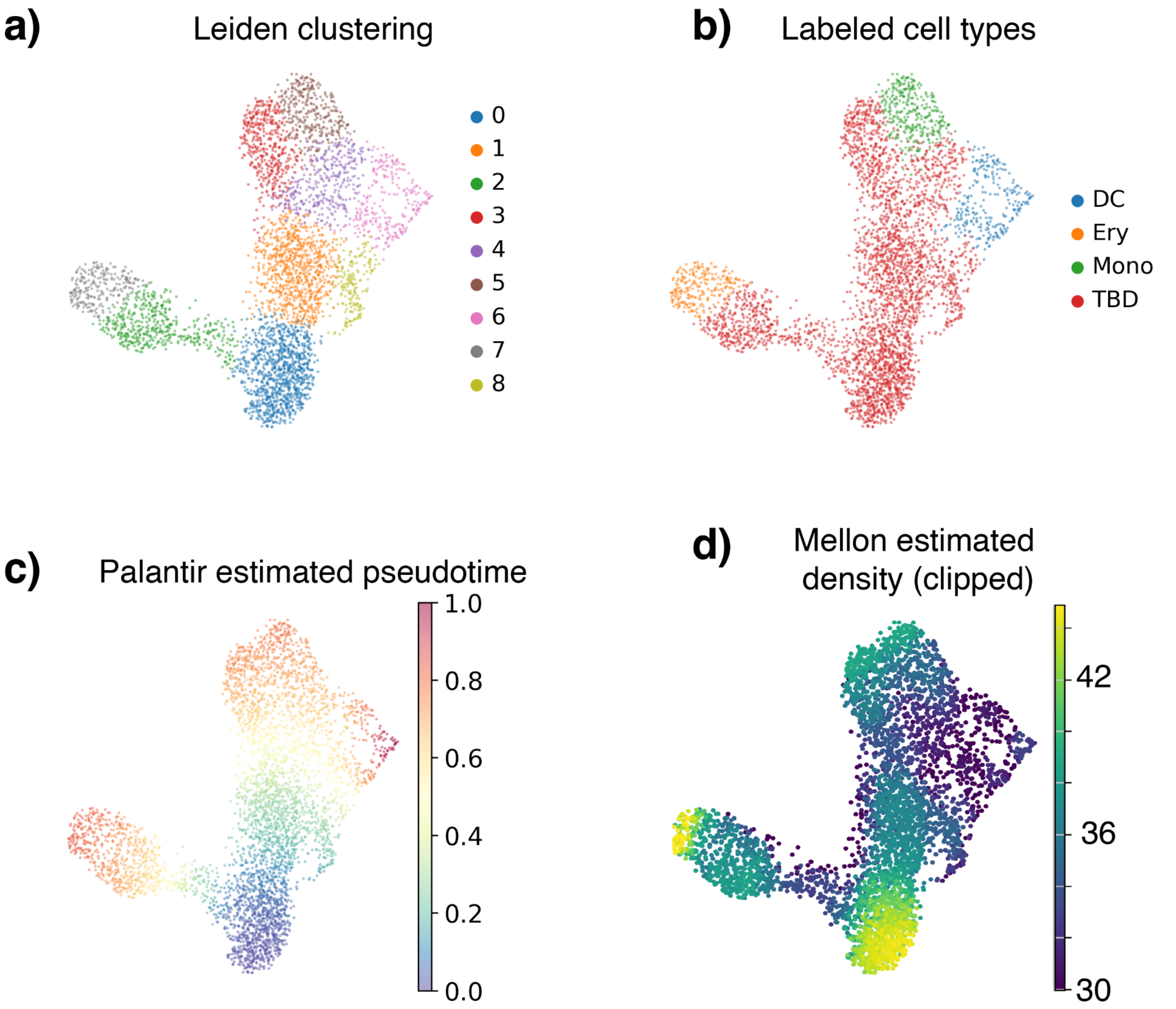


Figure S12: **Preprocess results for the Setty dataset.** **a)** Leiden clustering result using resolution 0.99 for the Setty dataset. **b)** UMAP plot showing the terminal cell types DC, Ery, and Mono. Monocytes correspond to cluster 5 in (a); Erythroids correspond to cluster 7; and DCs correspond to cluster 6. **c)** Pseudotime estimated by Palantir. **d)** Density (clipped) estimated by Mellon.

After we applied SCOPE, we plotted the size of the prediction set for all unlabeled cells, which displayed a smooth gradient of larger and larger prediction sets as the cell had a transcriptomic profile more dissimilar from the terminal cell types (Fig. S13a). A common diagnostic we found to be useful was tracking $\widehat{q}$ across recruitment iterations. Recall from the main text ("Method and analysis") that $\widehat{q}$ represents the $\alpha$ empirical quantile of the cumulative conformal scores. This means that a low $\widehat{q}$ qualitatively means that not many terminal cell types need to be included among the cells in the calibration set, which results in smaller prediction sets among the test (unlabeled) cells. As the recruitment iterations progress, we expected $\widehat{q}$ to gradually increase because not every semi-supervised label would be included among the top-predicted probabilities. (Note that $\widehat{q} = 1$ would necessarily yield a prediction set of all possible terminal cells.) Hence, we observed a gradual increase of $\widehat{q}$ after an initial dip in the first 10 recruitment iterations (Fig. S13b). This diagnostic reassured us that SCOPE's entropy balancing and adaptive conformal score were performing reasonably well, and that fate uncertainty propagated progressively through the SCOPE recruitment process from earlier to later cells. We next plotted all the initially unlabeled cells that had any of the three terminal cell types in their prediction sets (Fig. S13c-e). Lastly, we also performed the linear regression between entropy versus pseudotime to confirm that our estimated Mono-Ery branchpoint had a negative slope (Fig. S13f).

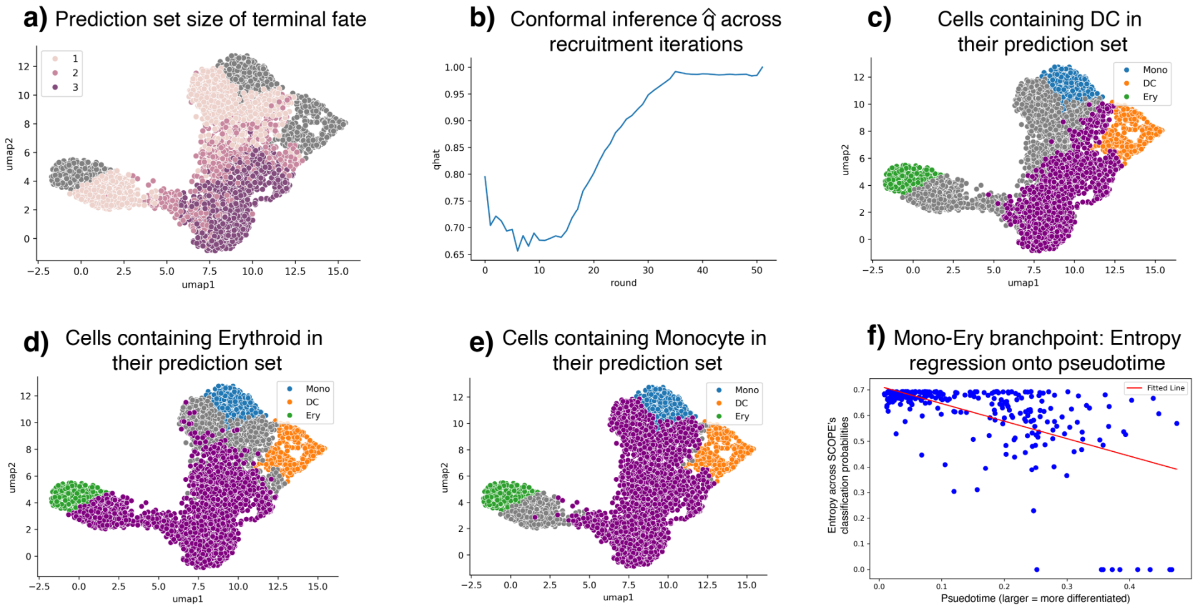


Figure S13: **Conformal results for the Setty dataset. a)** UMAP plot showing the size of the prediction set for the Setty dataset. **b)** The conformal score $\widehat{q}$ for each iteration. **c)** Initially unlabeled cells with DC in their prediction sets are highlighted in purple. **d)** Initially unlabeled cells with Ery in their prediction sets are highlighted in purple. **e)** Initially unlabeled cells with Mono in their prediction sets are highlighted in purple. **f)** Linear regression of entropy versus pseudotime for cells within the Ery-Mono branchpoint.

Next, we applied Slingshot [29] to the Setty dataset to assess if Slingshot could localize the branchpoint. We performed the experiment on this dataset, since Slingshot relies on an initial clustering of cells to determine principal curves that smoothly connect the cluster centers, prior to connecting the clusters using a minimum spanning tree (MST). (We found it difficult to perform this benchmark on the Weinreb dataset due to the Weinreb dataset's sheer size and complexity.) In principle, the cluster linking two diverging branches could be interpreted as a branchpoint. However, our analyses under three different Leiden clustering resolutions (Fig. S14-S16) show that Slingshot's inferred trajectories and the identity of the branchpoint cluster change markedly with the choice of resolution. At low resolution, Slingshot yields only coarse trajectories, while higher resolutions produce additional, often spurious, branches. This strong dependence on clustering granularity highlights a key limitation of such approaches: because their branchpoint definitions are tied to arbitrary clustering choices, they cannot reliably localize true biological branchpoints. Specifically, we first tried a Leiden clustering of resolution 0.25, where the resulting trajectories were reasonable, but the branchpoint between erythroid and the other two cell types was the entire red Cluster 0, and the branchpoint between monocytes and neutrophils was the entire yellow Cluster 1 (Fig. S15d). However, the larger resolution (i.e., larger clusters in Fig. S14) or smaller resolution (i.e., smaller clusters in Fig. S16) yielded vastly different results.

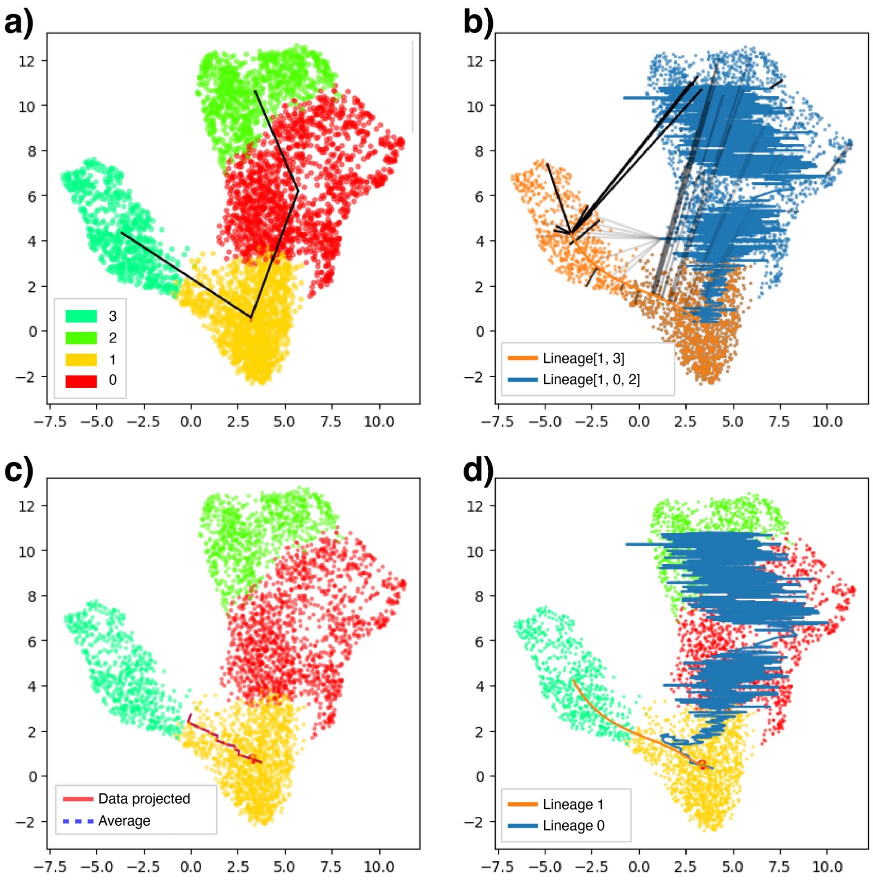


Figure S14: **Slingshot results for the Setty dataset using Leiden clustering at resolution 0.15. a)** Cluster-level minimum spanning tree (MST) constructed from the low-dimensional embedding, representing the global topology of cell-state transitions. **b)** Identification of lineage paths along the MST connecting the specified start node to terminal cell types (colored lines), along with the projection of a selection set of randomly chosen cells onto the lineage paths (black lines). **c)** Initialization of principal curves along the inferred lineage paths. **d)** Iteratively refined principal curves after trajectory fitting, representing the final inferred developmental trajectories. These lines are the same as those shown in (b).

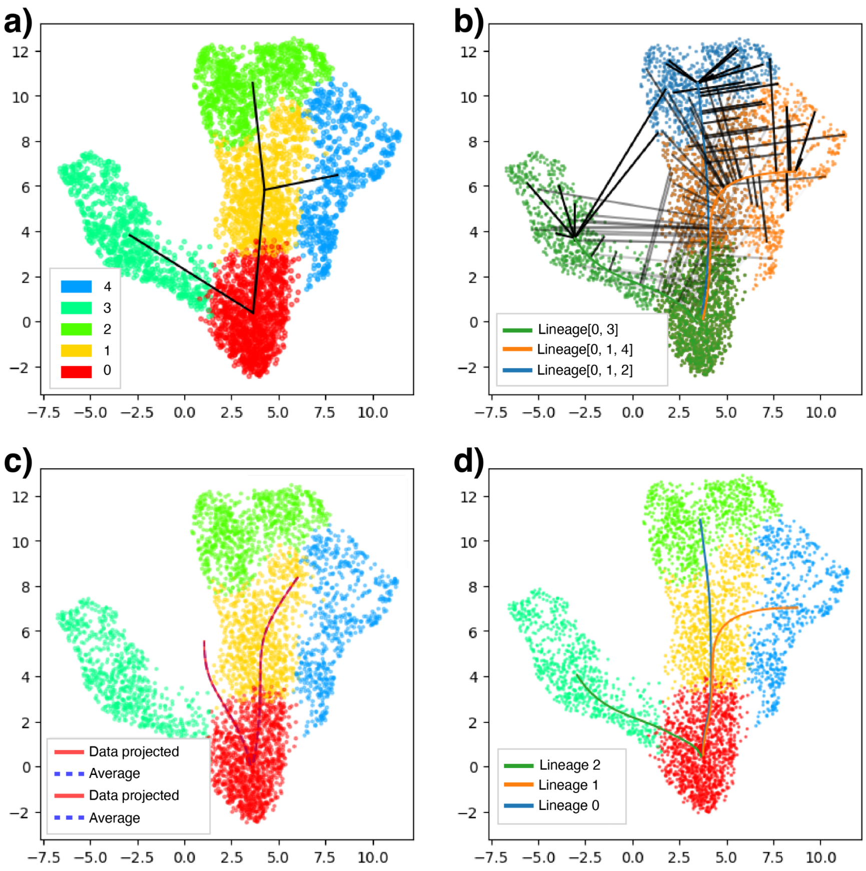


Figure S15: **Slingshot results for the Setty dataset using Leiden clustering at resolution 0.25.** The four plots mirror those shown in Fig. S14.

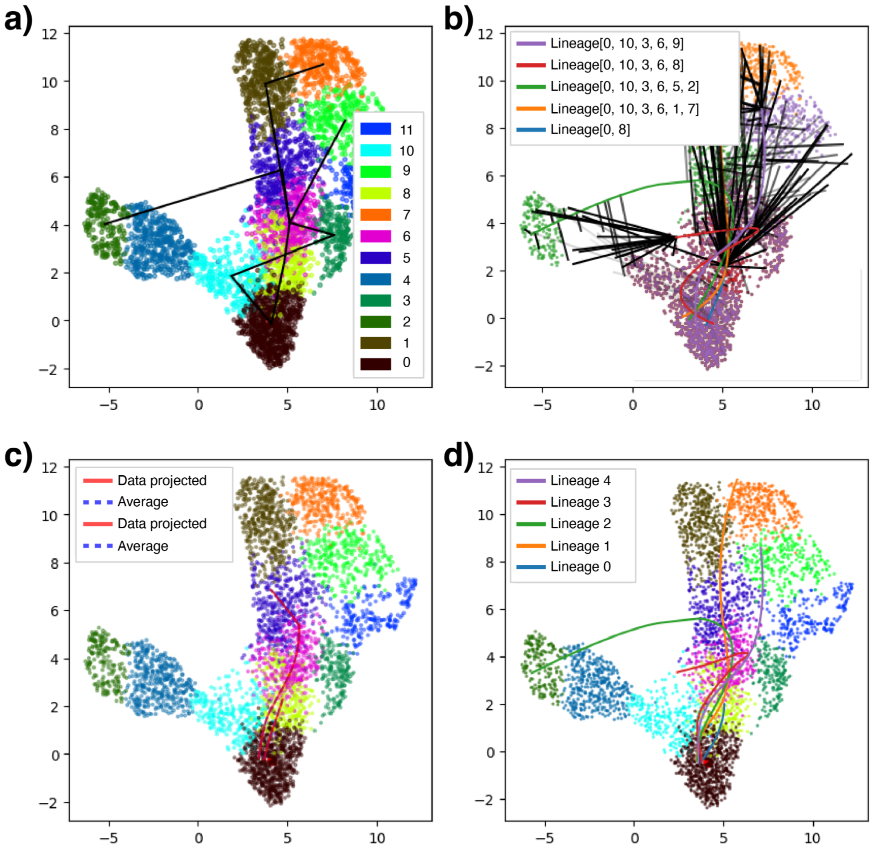


Figure S16: **Slingshot results for the Setty dataset using Leiden clustering at resolution 1.** The four plots mirror those shown in Fig. S14.

Next, we displayed plots about the preprocessing of the Persad dataset. In contrast to the Setty dataset, this dataset was provided with cell type labels (Fig. S17a). This yielded the estimated pseudotime (Fig. S17b). Additionally, when localizing branchpoints, SCOPE required the cell densities (Fig. S17c).

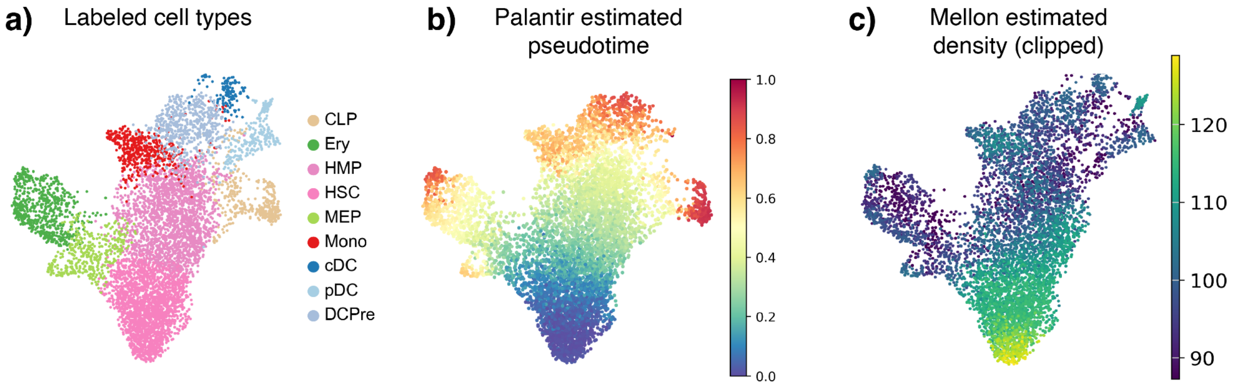


Figure S17: **Preprocess results for the Persad dataset. a)** UMAP plot showing provided cell type annotations (CLP, Common lymphoid progenitor; Ery, Erythroid; HSC, hematopoietic stem cell; HMP, hematopoietic multipotent progenitor; MEP, Megakaryocyte-erythroid progenitor; Mono, Monocyte; cDC, Conventional dendritic cell; pDC, Plasmacytoid dendritic cell; DCPre, Dendritic cell precursor). **b)** Pseudotime estimated by Palantir. **c)** Density (clipped) estimated by Mellon.

After we applied SCOPE, similar to Fig. S13, we plotted the size of the prediction set for all unlabeled cells, which also displayed a smooth gradient of larger and larger prediction sets as the cell had a transcriptomic profile more dissimilar from the terminal cell types (Fig. S18b). We also observed a gradual increase of $\widehat{q}$ after an initial dip in the first 10 recruitment iterations (Fig. S18b). We next plotted all the initially unlabeled cells that had any of the three terminal cell types in their prediction sets (Fig. S18c-e). Lastly, we also performed a linear regression of entropy versus pseudotime to confirm that our estimated Mono-Ery branchpoint had a negative slope (Fig. S18f).

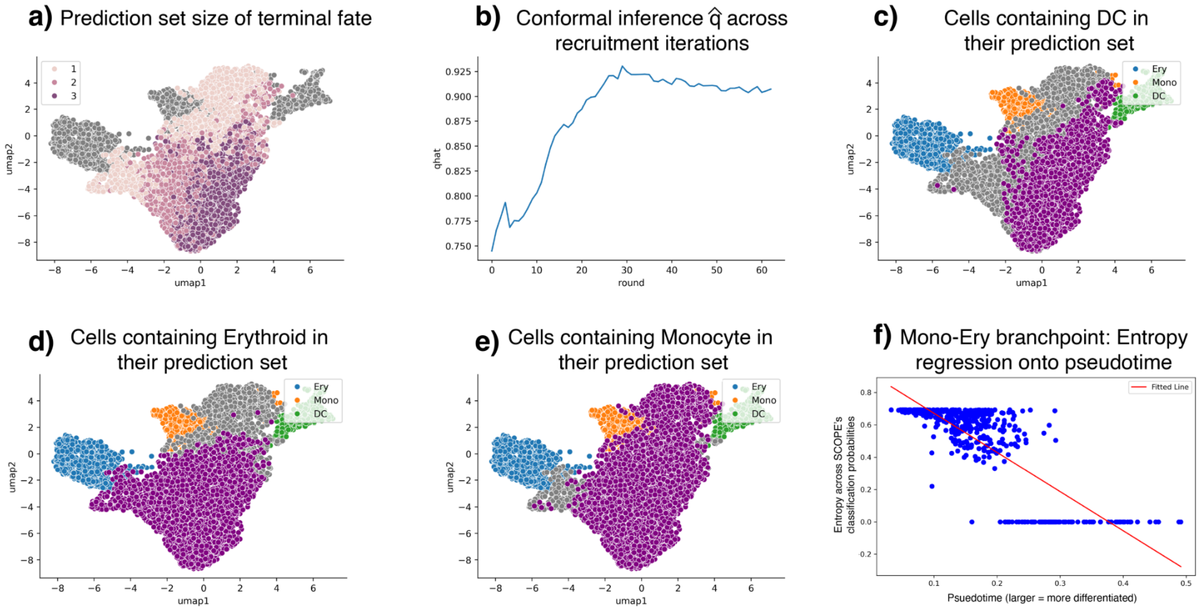


Figure S18: **Conformal results for the Persad dataset.** **a)** UMAP plot showing the size of the prediction set for the Persad dataset. **b)** The conformal score $\widehat{q}$ for each iteration. **c)** Initially unlabeled cells with DC in their prediction sets are highlighted in purple. **d)** Initially unlabeled cells with Ery in their prediction sets are highlighted in purple. **e)** Initially unlabeled cells with Mono in their prediction sets are highlighted in purple. **f)** Linear regression of entropy versus pseudotime for cells within the branchpoint of Ery and Mono.

After defining the Mono-Ery branchpoint in both the Setty and Persad datasets, we wanted to further confirm that the implicated TFs in our analyses (Fig. 4g-j) displayed the fate bias at the branchpoint that our importance score analyses suggested. Hence, in Fig. S19, we plot the gene expression of *CEBPD*, *CEBPA*, *NFIA*, *GATA2*, and *GATA1* among the cells localized in the Mono-Ery branchpoint in either dataset. Our UMAPs confirmed that cells with an elevated expression of *CEBPD* and *CEBPA* in the branchpoint had a fate bias towards the monocyte fate, while those with an elevated expression of *NFIA*, *GATA2*, and *GATA1* had a fate bias towards the erythroid fate.

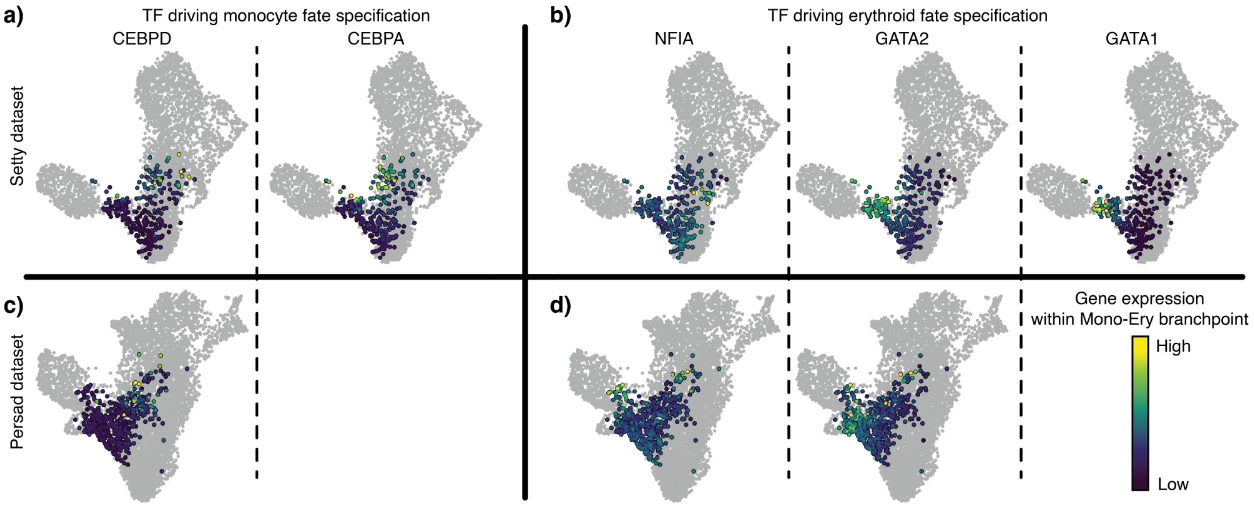


Figure S19: **Gene expression of driving TFs within Mono-Ery branchpoint a)** Gene expression of *CEPBD* and *CEPBA*, normalized within the cells in SCOPE's loalized branchpoint. **b)** Gene expression of *NFIA*, *GATA2*, and *GATA1*, normalized within the cells in SCOPE's loalized branchpoint. The UMAP coordinates are the same as in Fig. 4b-c. Not all genes are shown in the Persad dataset because they were not deemed highly variable.

Lastly, similar to Fig. S6, we investigated the number of cells with all combinations of terminal cell types in the conformal prediction sets in the Setty dataset (RNA), Persad dataset (RNA), and Persad dataset (ATAC) (Fig. S20a-c, respectively). All unique prediction sets across all three SCOPE analyses were shown. We noted strong similarities between the two RNA analyses: 1) the most common prediction set were the multipotent cells with all three terminal cell types in the prediction set, 2) the number of cells with {Mono, Ery} were larger than {Mono, DC}, suggesting that the first fate specification is the loss of potential to differentiate to either erythroids or DC cells, and 3) there were virtually no cells of prediction sets {Ery, DC}. These results supported our claim that SCOPE yielded reproducible results across two biological replicates. Furthermore, when comparing SCOPE's results between the RNA and ATAC modalities in the Persad dataset, we note that the number of singleton (i.e., unipotent) prediction sets was much higher in the ATAC modality. This further supported our findings shown in Fig. 5 of the main text, which showed that there are broad epigenetic priming mechanisms in the hematopoietic system.

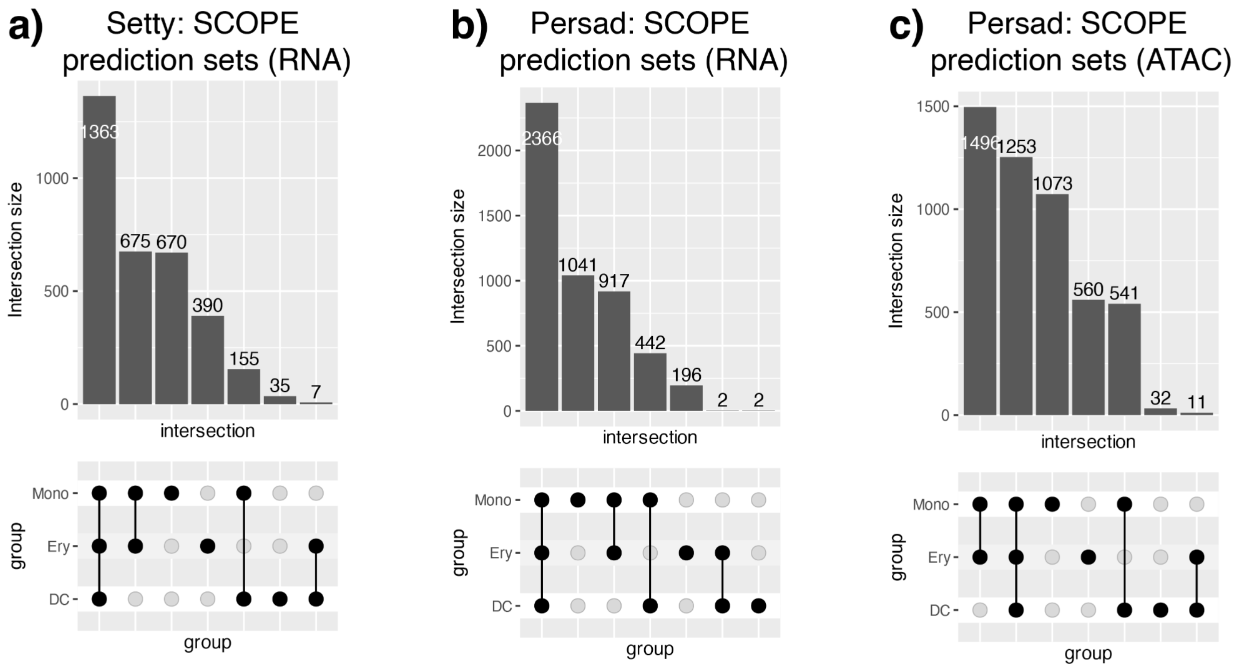


Figure S20: **Upset plot for the prediction sets among unlabeled cells in the Setty and Persad datasets. a)** SCOPE applied on the RNA modality of the Setty dataset. (Note: The Setty dataset only has the RNA modality.) **b)** SCOPE applied to the RNA modality of the Persad dataset. **c)** SCOPE applied to the ATAC modality of the Persad dataset.

#### S8.4 Additional results on Wohlschlegel dataset

In Fig. S21a-c, we showed the prediction sets for each cell, obtained by applying SCOPE to the ATAC modality, visualized on a UMAP derived from the ATAC modality. While almost all of the multipotent progenitor cells (MPCs) had the potential to differentiate to any of the three fates, as cells entered the (cycling) neurogenic precursors state (Npre and cyNpre), they started specifying their fates. In Fig. S21d, we visualize the CON-HRZ branchpoint when estimated from the ATAC modality.

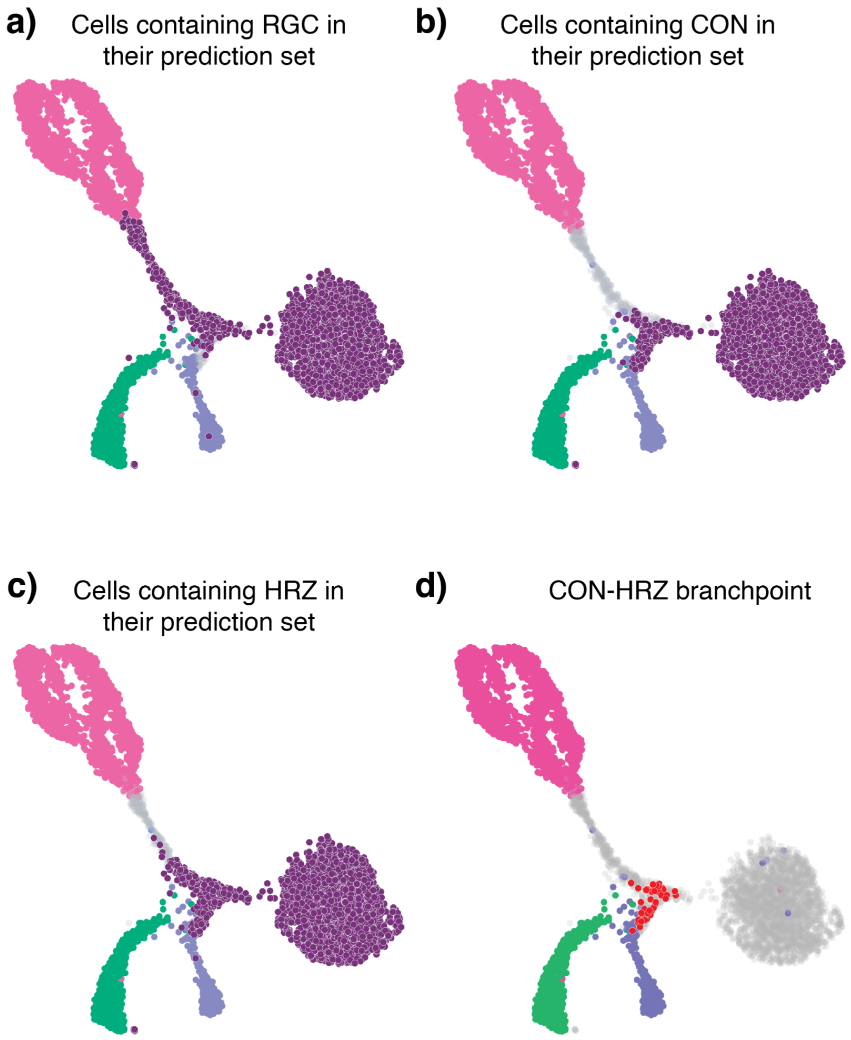


Figure S21: **Conformal results and branchpoint for the Wohlschlegel dataset. a)** Initially unlabeled cells with retinal ganglion cells (RGC) in their prediction sets are highlighted in purple. **b)** Initially unlabeled cells with cone photoreceptors (CON) in their prediction sets are highlighted in purple. **c)** Initially unlabeled cells with horizontal cells (HRZ) in their prediction sets are highlighted in purple. **d)** CON-HRZ branchpoint, highlighted in red. The UMAP and cell type coloring are retained from Fig. 5g. All results (both branchpoints and UMAP) reflected in this plot are derived from the ATAC modality.

Lastly, we systematically compared SCOPE with another method used to assess epigenetic priming, GrID-Net [41]. We highlighted four TFs relevant to cone photoreceptor differentiation: PRDM1, NR2E1, NEUROD1, and ONECUT2. Fig. S22a first visualized the importance scores of both `PRMD1` and `NR2E1` (red) and their corresponding aggregated enhancers, both of which were implicated as two of the six TFs displaying epigenetic priming in Fig. 5l. Below are the results of GrID-Net, analyzed via `gridnet.run_gridnet()`, after providing the method with the same set of linked peak-genes. Briefly, GrID-Net outputted two types of results relevant to our analysis: (1) the normalized expression of gene expression and aggregated enhancers across pseudotime, and (2) the number of linked enhancers deemed to display priming (predictive) via Granger causality (i.e., changes in the enhancer accessibility forecast changes in the gene expression). We noted some distinctions between SCOPE and GrID-Net: While both methods indicate that enhancer accessibility reaches its peak prior to gene expression, the temporal difference is quite subtle. Furthermore, GrID-Net was unable to implicate any enhancers individually due to the small signal size. In contrast, SCOPE uses all the enhancers collectively to predict the cone photoreceptor fate, and afterwards investigates the importance of the linked enhancers to a particular TF after fitting the classifier. This workflow allowed for a more granular appreciation of epigenetic priming.

Fig. S22b showed SCOPE's and GrID-Net's results for NEUROD1 and ONECUT2 as a contrast. Neither SCOPE nor GrID-Net could determine epigenetic priming. However, recall that SCOPE not finding evidence for epigenetic priming (and, related, the MPCs having a prediction set of all three fates) was not necessarily a claim that the MPCs have not specified their fate for cone photoreceptors. This is for two reasons: (1) Since the Wohlschlegel dataset is only of Day 59 cells, the lack of epigenetic priming is not necessarily "proof" of no priming, (2) in low-power settings, SCOPE's prediction sets naturally cover all possible fates, meaning no priming would be detected.

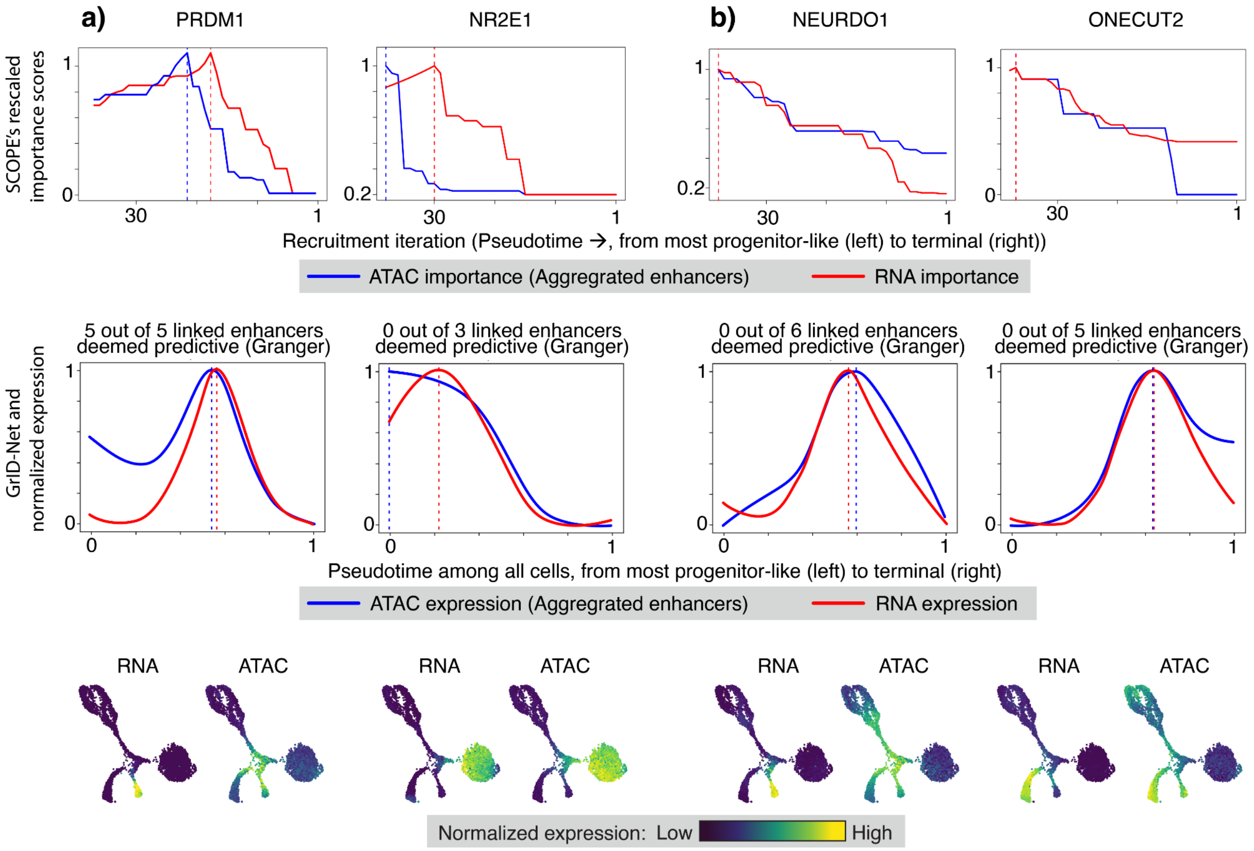


Figure S22: **Epigenetic priming using SCOPE and GrID-Net.** **a)** Top row: SCOPE's normalized importance scores of PRDM1 and NR2E1, both TFs implicated to have epigenetic priming by SCOPE. The plots are in the same layout as Fig. 5j, where the x-axis shows the recruitment iteration (later iterations: more progenitor-like cells). The apexes of the aggregated enhancers (blue) and the gene (red) are marked with dotted horizontal lines. Middle row: GrID-Net's results, showing the aggregated enhancer accessibility and gene expression (normalized) against pseudotime, with the apex also marked. Bottom row: UMAPs of the TFs, either based on the gene expression (RNA) or aggregated enhancer accessibility (ATAC). **b)** Same format as (a), but for NEUROD1 and ONECUT2, two other TFs that are known to drive cone photoreceptor differentiation.

[22] Andrew Ng, Michael Jordan, and Yair Weiss. On spectral clustering: Analysis and an algorithm. *Advances in Neural Information Processing Systems*, 14, 2001.

[31] Ryan J Tibshirani, Rina Foygel Barber, Emmanuel Candes, and Aaditya Ramdas. Conformal prediction under covariate shift. *Advances in neural information processing systems*, 32, 2019.

[32] Cole Trapnell, Davide Cacchiarelli, Jonna Grimsby, Prapti Pokharel, Shuqiang Li, Michael Morse, Niall J Lennon, Kenneth J Livak, Tarjei S Mikkelsen, and John L Rinn. Pseudo-temporal ordering of individual cells reveals dynamics and regulators of cell fate decisions. *Nature Biotechnology*, 32(4):381, 2014.

[33] Vladimir Vovk, Alexander Gammerman, and Glenn Shafer. *Algorithmic Learning in a Random World*. Springer, Cham, 2 edition, 2005.
